# Lost and found: *beer belly* is a novel, ancient regulator of animal development

**DOI:** 10.64898/2026.10.24.761492

**Authors:** Zoë X. Schultz, Grace Blakeley, Madeleine E. Aase-Remedios, Dorothea V. Grillmayer, Daniel J. Leite, Tom Frankish, Gemma Sutton, Grigory Genikhovich, Steffen Scholpp, Matthias Pechmann, José M Martín-Durán, Natascha Turetzek, Ralf Janssen, Alistair P. McGregor

## Abstract

Single-cell RNA sequencing (scRNA-seq) can provide new insights into the regulation of developmental processes by revealing unknown or overlooked genes. We identified a novel gene called *beer belly* (*bebe*) as a marker of a segment addition zone (SAZ) associated cell cluster in a single-cell atlas of the spider *Parasteatoda tepidariorum*. Here we show that *bebe* is an ancient toolkit gene encoding a previously uncharacterised protein predicted to be secreted, which likely plays an important role in the development of most animal phyla. However, we could not identify a *bebe* ortholog in any mammal or holometabolous insect, suggesting it has been lost in these lineages and explaining why it has been overlooked to date. Knock-down of *bebe* expression in *P. tepidariorum* demonstrates that it regulates SAZ size and consequently segment size by interacting with *Wnt8*. Our findings also suggest that *bebe* may regulate segmentation in other invertebrates and play additional developmental roles in bilaterian and non-bilaterian animals, and likely ancestrally, through modulation of Wnt signalling. This study emphasises the promise of applying single-cell sequencing across a diverse range of species to better understand the regulation of development beyond candidate genes from traditional models, allowing us to better characterise the animal developmental toolkit and its evolution.

## Introduction

Single-cell RNA sequencing (scRNA-seq) can provide new insights into development because it has the potential to reveal unknown or overlooked genes involved in the regulation of developmental processes^1,2^. Fulfilling this great potential requires interrogation of the function of new marker genes of key cell types in developing tissues identified in unbiased scRNA-seq screens and how they interact with previously known candidate genes and pathways.

Panarthropods, annelids and chordates are the three major animal groups with segmented bodies. These metameric body plans allow the specialisation of individual segments, which is thought to have contributed to the evolutionary success of these species-rich and highly morphologically diverse lineages^3^. Although the evolutionary origins of segmentation are debated, it is probable that this process evolved independently at least between protostomes and deuterostomes^3-8^. It is also well supported that the last common ancestor of arthropods was segmented and generated posterior segments sequentially from a growth zone or segment addition zone (SAZ), as seen in most extant arthropods^4,9,10^.

Studies in a range of arthropods over the last few decades mainly using candidate gene approaches have revealed conserved and divergent features of the regulation of sequential segment addition (reviewed in ^9,10,11,12^). In the ancestral arthropod, posterior segment addition was likely regulated by upstream Wnt and Delta-Notch (*Dl-N*) signalling in the SAZ, directing dynamic expression of downstream genes, including *caudal* (*cad*) and pair-rule gene orthologues, in a clock-like mechanism to specify the expression of segment polarity genes and segmental boundaries (e.g. ^10,11,13-20^). However, we still have a very limited understanding of SAZ formation and functionality, and in particular how segment size is specified. Despite the success of the candidate gene approach, it has focused on previously known animal developmental ‘toolkit’ genes and relied on only a few model organisms, including the fruit fly *Drosophila melanogaster*, which generates its segments almost simultaneously without a SAZ. Therefore, unbiased approaches across a range of species with sequential segment addition, as in the arthropod ancestor, are needed to better understand the genetic regulation of segment addition and how it evolved among arthropods.

The spider *Parasteatoda tepidariorum* (formerly *Achaearanea tepidariorum*) serves as a key chelicerate model for investigating segmentation and other aspects of development, particularly as a point of comparison to other arthropod models in the Mandibulata^21,22^. *P. tepidariorum* generates its twelve opisthosomal segments, which comprise the tagma posterior to the leg-bearing cephalothorax, sequentially from a posterior SAZ. The SAZ is specified before germband formation by expression of the ligand-encoding genes *Wnt8* and *Delta* (*Dl*), and RNAi knockdown of either of these genes results in truncated germbands often lacking all opisthosomal segments^15,23^. Wnt8 and Delta-Notch signalling combine to regulate the dynamic expression of *caudal* (*cad*), *even skipped* (*eve*) and other segmentation genes in the SAZ^15,23,24^. *Wnt8* likely maintains a pool of cells in the posterior SAZ, but suppression of *Wnt8* by *Dl* in anterior cells allows the formation of nascent segments marked by stripes of expression of genes including *eve* and *cad*. These subsequently regulate expression of the segment polarity gene *engrailed* (*en*) and thus formation of segmental boundaries^15,24^.

To understand the regulation of segmentation further beyond known candidate genes, we previously carried out scRNA-seq of embryos of the spider *P. tepidariorum* during developmental stages when segmentation occurs^25^. In our scRNA-seq atlas, we identified a cluster of cells marked by known SAZ genes, including *Wnt8*, *cad*, and *eve*^25^. In addition, one of the most significant marker genes of the SAZ cluster was *g30822* (aug3.g27670^26^, LOC107442497), hereafter called *beer belly* (*bebe*)^25^. This gene is expressed in posterior SAZ cells overlapping with *Wnt8* expression and responds to Hedgehog (Hh) signalling^25,27^. However, the evolutionary history, developmental role of this novel SAZ marker, and its contribution to spider segmentation remained unknown.

Here we combine broad phylogenetic searches and functional approaches in *P. tepidariorum* to characterise the evolution, expression and function of *bebe*. We found *bebe* orthologues in many animal phyla, suggesting it is an ancient animal gene encoding a previously uncharacterised protein predicted to be secreted. However, this gene has an unusual phylogenetic distribution because *bebe* has likely been lost in several animal lineages, including mammal, nematode, annelid and insect model organisms. We also analysed the expression of *bebe* further in *P. tepidariorum* and characterised the expression of *bebe* orthologues in a range of other bilaterians and a cnidarian. This suggests that the function of *bebe* is potentially conserved in other groups of segmented invertebrates and that this gene plays important roles during embryogenesis in other bilaterian and non-bilaterian animals and, therefore, likely ancestrally. We then demonstrate that in *P. tepidariorum*, *bebe* plays a role in specifying SAZ size and consequently segment size by interacting with *Wnt8* signalling. Taken together we have identified a novel but ancient animal development toolkit gene that may modulate Wnt signalling to regulate developmental processes across animals.

## Results

### *bebe* is an ancient and common gene but has likely been lost in several animal lineages

To explore the evolution of *bebe* we searched for its orthologues using BLAST searches across taxonomic subsets of the GenBank database. We found that *bebe* is shared across representatives of all major groups of bilaterians and cnidarians but was likely lost several times independently in various animal lineages (Fig. 1 and Fig. S1, Supplementary File 1). *bebe* is present in other spiders and scorpions but not harvestmen, only five orders of insects (Zygentoma, Entomobryomorpha, Ephemeroptera, Thysanoptera, and Hemiptera), most mollusc orders, some annelids but not the model species *Platynereis dumerilii*, and several fish orders. Furthermore, in lineages that underwent whole-genome duplications (WGDs), including arachnopulmonates, vertebrates, and teleosts, *bebe* is rarely retained in duplicate, except in cases of multiple successive and recent WGDs (i.e. we found two copies in horseshoe crabs and two copies in a carp) (Fig. S1, Supplementary File 1). However, we found no *bebe* orthologues in holometabolous insects, including the model species *D. melanogaster* or *Tribolium castaneum*, amphibians, birds, mammals, or non-metazoan eukaryotes. (Fig. 1 and Fig. S1, Supplementary File 1). While some of the apparent absences of *bebe* across the animal phylogeny may be due to a lack of high-quality reference genomic data, we are more confident in the absences from model organisms for which multiple data sources are available (e.g., multiple genome assemblies and transcriptomes). Furthermore, entries on OrthoDB (group 986800at33208) and PANTHER (PTHR35842) are consistent with the phylogenetic distribution we observed. Outside animals, we identified a few bacterial sequences with low similarity to part of the Bebe protein sequence, but these bacterial genes are predicted to encode a domain with peptidase function not found in any Bebe orthologues, and therefore we consider that it is unlikely they are related to animal Bebe (Supplementary File 1).

**Figure 1.**
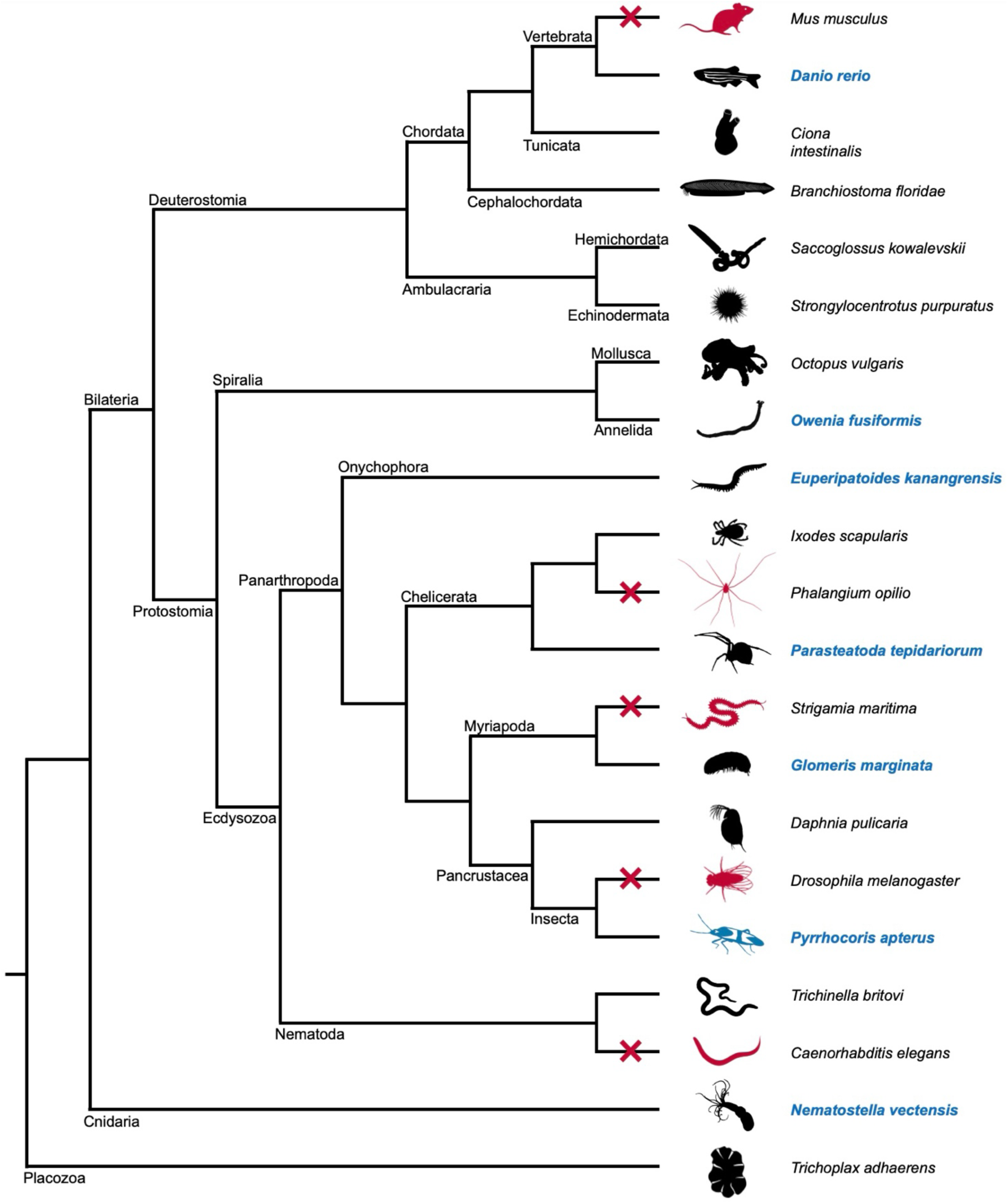
Summary of the phylogenetic distribution of *bebe*. Animal phylogeny showing the distribution of *bebe* orthologues. Animals indicated in red are in lineages that may have lost *bebe* (red crosses). Animals shown in black have retained *bebe* with those named in blue font surveyed for *bebe* expression during embryogenesis (Figs 2 and 3 and Figs S4-12). Images were sourced from PhyloPic under the following licences: *M. musculus*, *D. rerio*, *C*. *intestinalis*, *B. floridae*, *S. kowalevskii*, *S. purpuratus*, *O. vulgaris*, *E. kanangrensis*, *P. opilio*, *S. maritima*, *G. marginata*, *D. pulicaria*, *D. melanogaster*, *P. apterus*, *T. britovi*, *D. elegans*, *N*. *vectensis and T. adhaerens* (https://creativecommons.org/publicdomain/zero/1.0/); *O. fusiformis* by Anne Weigert and Christoph Bleidorn (https://creativecommons.org/licenses/by-nc-sa/3.0/); *I. scapularis* and *P. tepidariorum* by Mathilde Cordellier (https://creativecommons.org/licenses/by-nc/3.0/).

None of the identified *bebe* orthologues was annotated with a name or function in the searched genomic databases and so we next studied the predicted structure of Bebe to infer more about the function of this protein. The Bebe orthologues of a vertebrate and several invertebrates, including *P. tepidariorum*, are predicted to have similar structures with two nearly parallel beta-sheets, but no other conserved domains with known functions were indicated that might reveal what type of protein *bebe* encodes (Fig. S2). However, we detected a predicted N-terminal signal peptide in most surveyed orthologues, suggesting Bebe might be secreted (Fig. S3).

### *bebe* is expressed in the posterior SAZ in the spider *Parasteatoda tepidariorum*

It was previously shown that *Pt-bebe* marks SAZ cell clusters and is expressed in the posterior cells of this tissue during segmentation as well as in the head and differentiating segments^25,27,28^. We show here that *Pt-bebe* is expressed at stage 6, as the embryo transitions from radial to axial symmetry, in a small circular posterior domain in the region of the blastopore in the forming SAZ (Fig. 2A). At stage 7, *Pt-bebe* is expressed broadly throughout the SAZ and in a nascent stripe at the anterior edge of the germband (Fig. 2B). At stage 8.1, strong expression persists in the SAZ, and the anterior stripe has moved posteriorly away from the rim of the germband (Fig. 2C). At this stage expression appears in the ventral region of the cheliceral, pedipalpal, and leg-bearing segments (Fig. 2C,D), which develop into stripes at stage 8.2. The head stripe subsequently splits into a separate domain in each head lobe and expression is still observed in the ventral prosomal segments as well as in the posterior SAZ at stage 9.1 (Fig. 2E). Similar expression is seen at stage 9.2 (Fig. 2F). At stage 10.1 expression is visible in the prosomal appendages and the opisthosomal limb buds in segments O2 to O5 and expression appears in the stomodaeum, while the head lobe expression contracts into smaller domains (Fig. 2G). Interestingly, the posterior SAZ domain becomes thinner from stage 7 to stage 10.1 as development progresses and opisthosomal segments are added (Fig. 2B-G).

**Figure 2.**
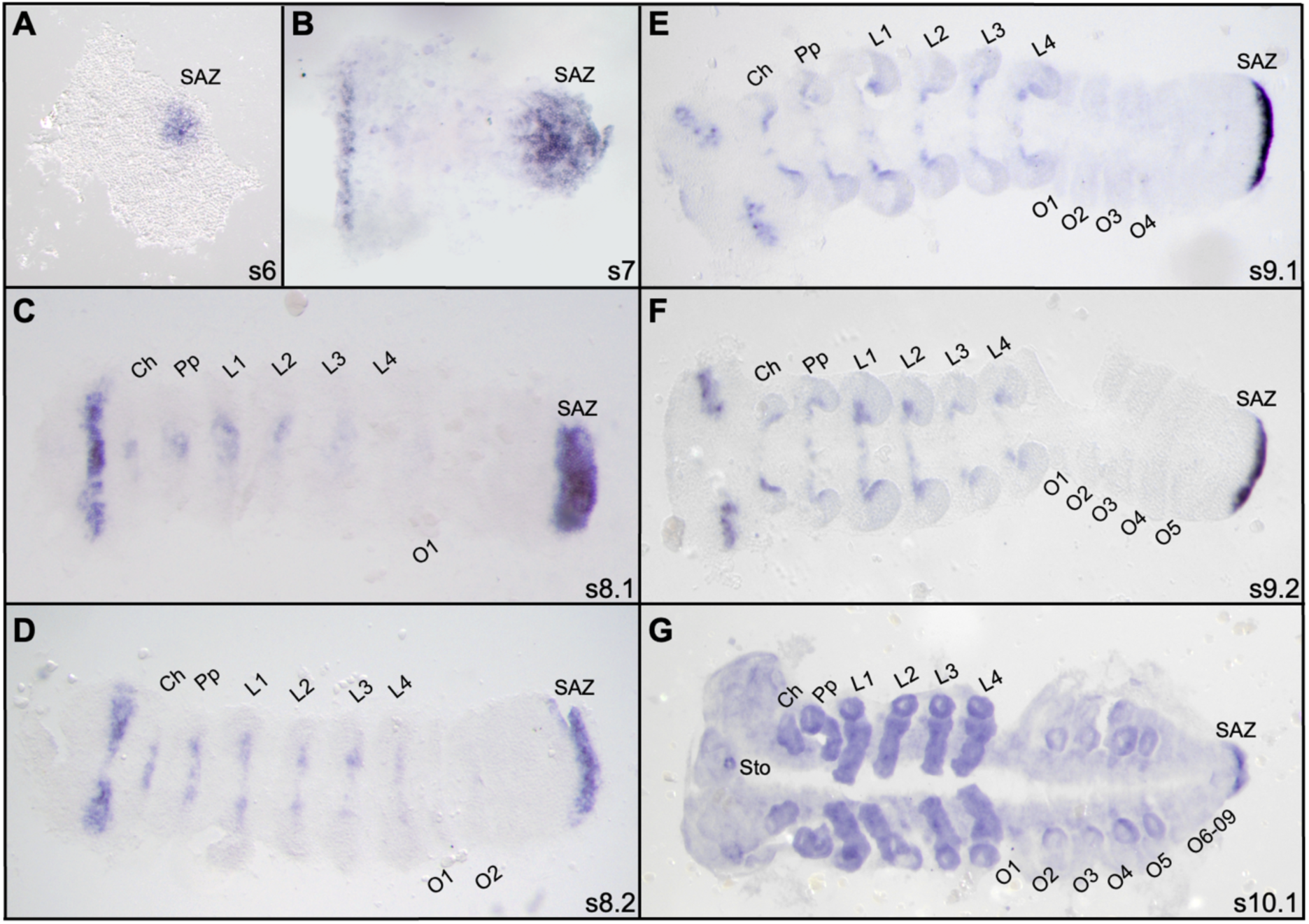
*bebe* expression in *P. tepidariorum* embryos. *bebe* is expressed from stage 6 in the SAZ (A) with a stripe forming in the head at stage 7 (B). Expression persists in the SAZ as segments are added through stages 7 to 10.1 (B-G). Stripes of expression start to form from stage 8.1 in the ventral region of prosomal segments (C). The head stripe begins to split into two separate domains from stage 8.2. At stages 9.1 and 9.2 expression persists in a domain in each head lobe, in the ventral of prosomal segments and in the SAZ (E and F). Expression is seen around the stomodeum and in the developing prosomal appendages and opisthosomal limb buds at stage 10.1 (G). Sto, stomodeum; Ch, cheliceral segment; Pp, pedipalpal segment; L, leg-bearing segment; O, opisthosomal segment; SAZ, segment addition zone. Embryos are flat-mounted with the anterior to the left.

### *bebe* expression during embryogenesis in other species

To compare *bebe* expression in *P. tepidariorum* with other animals, we surveyed the expression of *bebe* orthologues in several species at successive phylogenetic distances to this spider, including a wolf spider *Pardosa amentata*, the cellar spider *Pholcus phalangioides*, the tarantula *Acanthoscurria geniculata*, the millipede *Glomeris marginata*, the hemipteran insect *Pyrrhocoris apterus*, the onychophoran (velvet worm) *Euperipatoides kanangrensis*, the annelid *Owenia fusiformis*, the zebrafish *Danio rerio*, and the cnidarian *Nematostella vectensis* (Fig. 3 and Figs S4 to S12).

**Figure 3.**
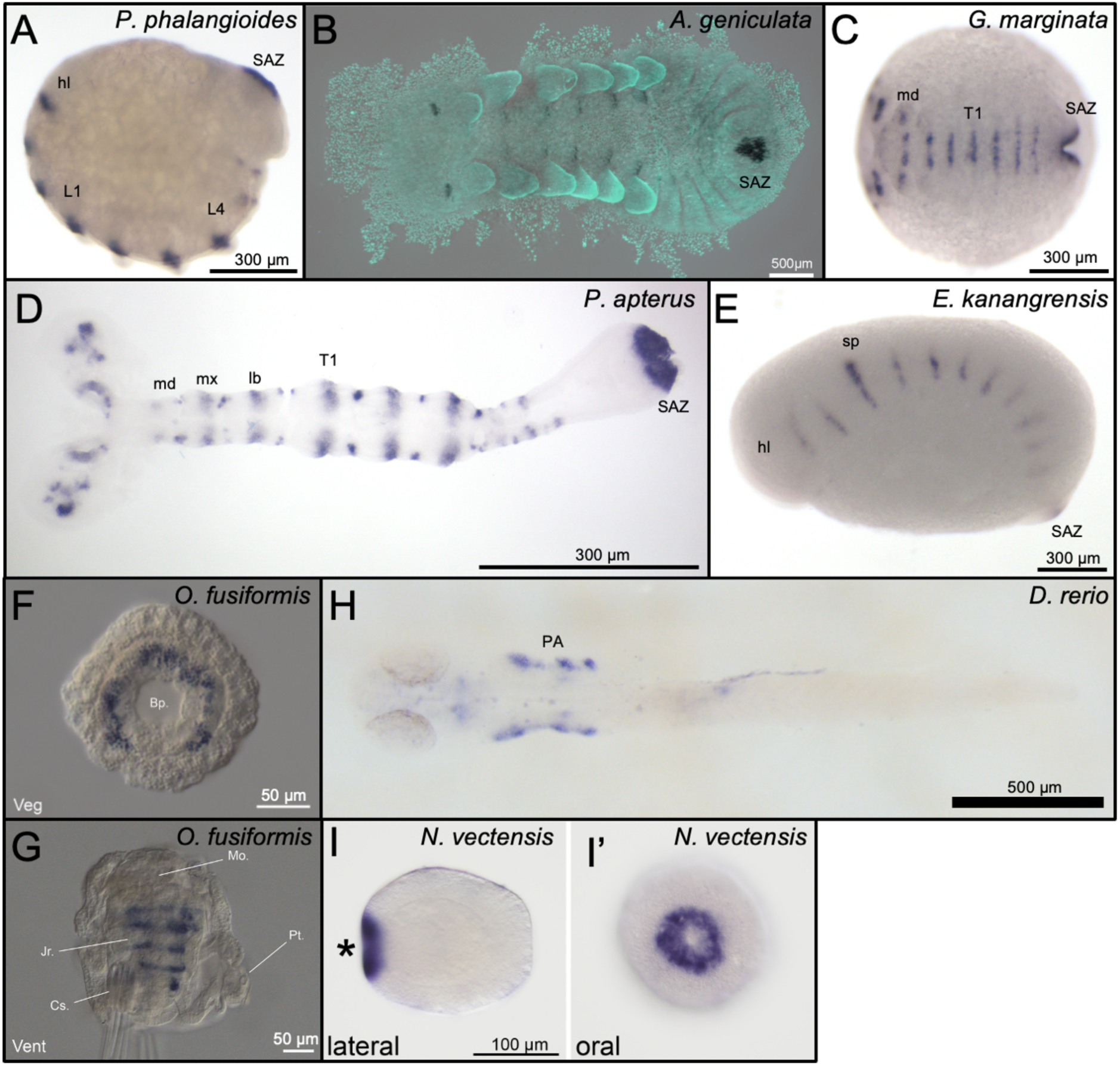
*bebe* expression in other selected animals. In the germband stage embryos of the spiders *P. phalangioides* (A) and *A. geniculata* (B), *bebe* is expressed in the head lobes (hl), ventral region of the prosomal segments and SAZ (L1 and L4 = leg bearing segments 1 and 4). In *G. marginata* embryos, *bebe* is expressed in the head and segmental stripes as well as in the SAZ (T1 = first trunk segment) (C). In *P. apterus* extending germband stage embryos, *bebe* is expressed in the developing head lobes, around the stomodeum, segmentally in the gnathal (md = mandibular, mx = maxillary, lb = labial), thoracic (T1) and anterior abdominal segments as well as the SAZ (D). In *E. kanangrensis*, *bebe* expressed in segmental stripes and in a domain at the posterior of the germband (hl = head lobe, sp = slime papilla) (E). In the gastrula of *O. fusiformis, bebe* is expressed in the endomesoderm, forming an incomplete ring of expression around the blastopore (Bp) (F). In *O. fusiformis* larvae at ∼21 dpf, *bebe* is expressed in lateral stripes at the boundaries of developing segments in the juvenile trunk rudiment. Expression is also detected in the buccal organ and the brain. Mo = mouth; Pt = prototroch; Jr = juvenile rudiment; Cs = chaeta-sac (G). In *D. rerio* embryos at 24 hpf, *bebe* is expressed in the pharyngeal arches (PA) and tegmentum (flat-mounted dorsal view with anterior to the left) (H). In *N. vectensis* early planula larva at 2 days post-fertilization, *bebe* transcripts are present in the blastopore (*) (I and I’).

The expression of *bebe* orthologues is generally conserved among the investigated arthropods (Fig. 3 and Figs S4 to S8). In addition, segmentally reiterated expression and expression in the SAZ is also evident in the onychophoran (Fig. 3 and Fig. S9). In the annelid, *bebe* is expressed asymmetrically around the blastoporal lip, potentially in the developing mesoderm in gastrula embryos and later in segmental stripes (Fig. 3 and Fig. S10). In the zebrafish, *D. rerio*, *bebe* is first expressed in presumptive placodal cells and later in the pharyngeal arches and tegmentum (Fig. 3 and Fig. S11). Finally in the anthozoan cnidarian *N. vectensis, bebe* is expressed around the blastopore (Fig. 3 and Fig. S12). Taken together, these results suggest *bebe* is involved in segmentation in other segmented invertebrates, as well as spiders, and *bebe* orthologues likely play several other roles during embryogenesis in bilaterians and non-bilaterians.

### Knockdown of *Pt-bebe* results in enlargement of the developing opisthosoma

To study the function of *bebe* during spider embryogenesis we used parental RNAi (pRNAi) to knock down the expression of this gene in *P. tepidariorum* (Fig. 4, Figs S13 and S14). At stage 7, the developing opisthosoma, including the SAZ, of *Pt-bebe* pRNAi embryos is wider along the D-V axis compared to controls and the surface of the germband appears uneven (Fig. 4A-B’). Subsequently during stage 8.1 and 8.2, the more posterior opisthosomal segments are wider and appear distorted, although *Pt-bebe* pRNAi embryos can generate segments O1-O3 with relatively normal morphology (Fig. 4C-D’). The SAZ is also wider than normal at these stages (Fig. 4C-D’). As further segments are added during stages 9.1 to 10.1, they also appear wider than normal and distorted, and the SAZ appears perturbed and does not become narrower like in control embryos (Figs 4E-H’ and 5). We did not observe any *Pt-bebe* pRNAi embryos with more than eight or nine opisthosomal segments even in milder knockdowns, indicating a potential failure-point in normal development as the developing head and opisthosoma abut (Fig. S15). In some stage 9.1 to 10.1 *Pt-bebe* pRNAi embryos, we also observe over-growth of head lobes and distortion of this tissue, but the appendage-bearing prosomal segments appear to develop normally (Fig. 4 and Fig. S14). Taken together, these results show that, consistent with *Pt-bebe* expression patterns, this gene is required for correct growth of the head and opisthosoma, and although most posterior segments can still form when *Pt-bebe* is knocked-down, they are larger and appear distorted.

**Figure 4.**
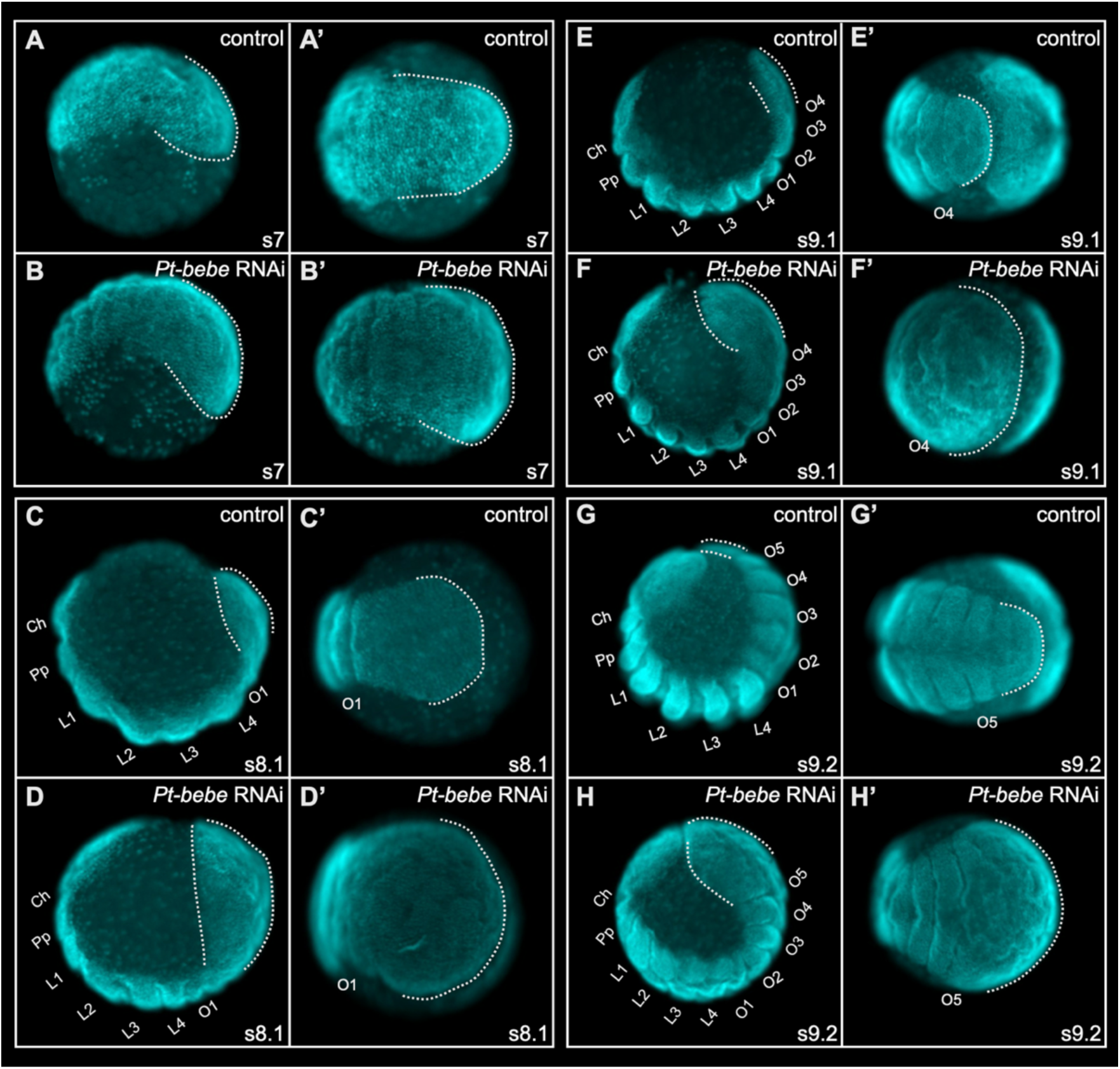
Parental RNAi knockdown of *Pt-bebe*. Control and *Pt-bebe* RNAi embryos at stages 7 (A-B’), 8.1 (C-D’), 9.1 (E-F’) and 9.2 (G-G’). Lateral views with anterior to the left and ventral down (A-D and E-H) and ventral views of the opisthosoma with anterior to the left (A’-D’ and E’-H’). Dotted lines indicate the posterior opisthosoma including the SAZ. Ch, cheliceral segment; Pp, pedipalpal segment; L, leg-bearing segment; O, opisthosomal segment. All embryos are stained with DAPI.

### Segment boundaries form normally, but the segments contain more cells upon *Pt-bebe* RNAi

We then investigated whether segment boundaries still form normally when *Pt-bebe* is knocked-down by assaying the expression of *Pt-en-1*, a marker of posterior segmental boundaries (Fig. 5 and Fig. S16)^29^. In whole-mount *Pt-bebe* pRNAi embryos with distorted opisthosomas *Pt-en-1* expression is still observed in the segments (Fig. S16). When we flat-mounted these embryos *Pt-en-1* expression appeared in regular, intact stripes at the posterior boundary of each formed segment (Fig. 5). However, in these embryos, nascent *Pt-en-1* stripes appeared farther apart along the A-P axis, and most opisthosomal segments appeared wider along the D-V axis than in controls (Figs 4 and 5). This suggests that while *Pt-bebe* is not required for formation of posterior segments *per se*, knockdown of this gene does affect segment size. To test this further, we examined the expression of the segmentation gene *Pt-eve* and found it is expressed in a similar pattern in *Pt-bebe* pRNAi and wild-type embryos, although the stripes of expression were longer along the D-V axis in the knockdown embryos, consistent with the wider opisthosomal segments (Fig. S17).

**Figure 5.**
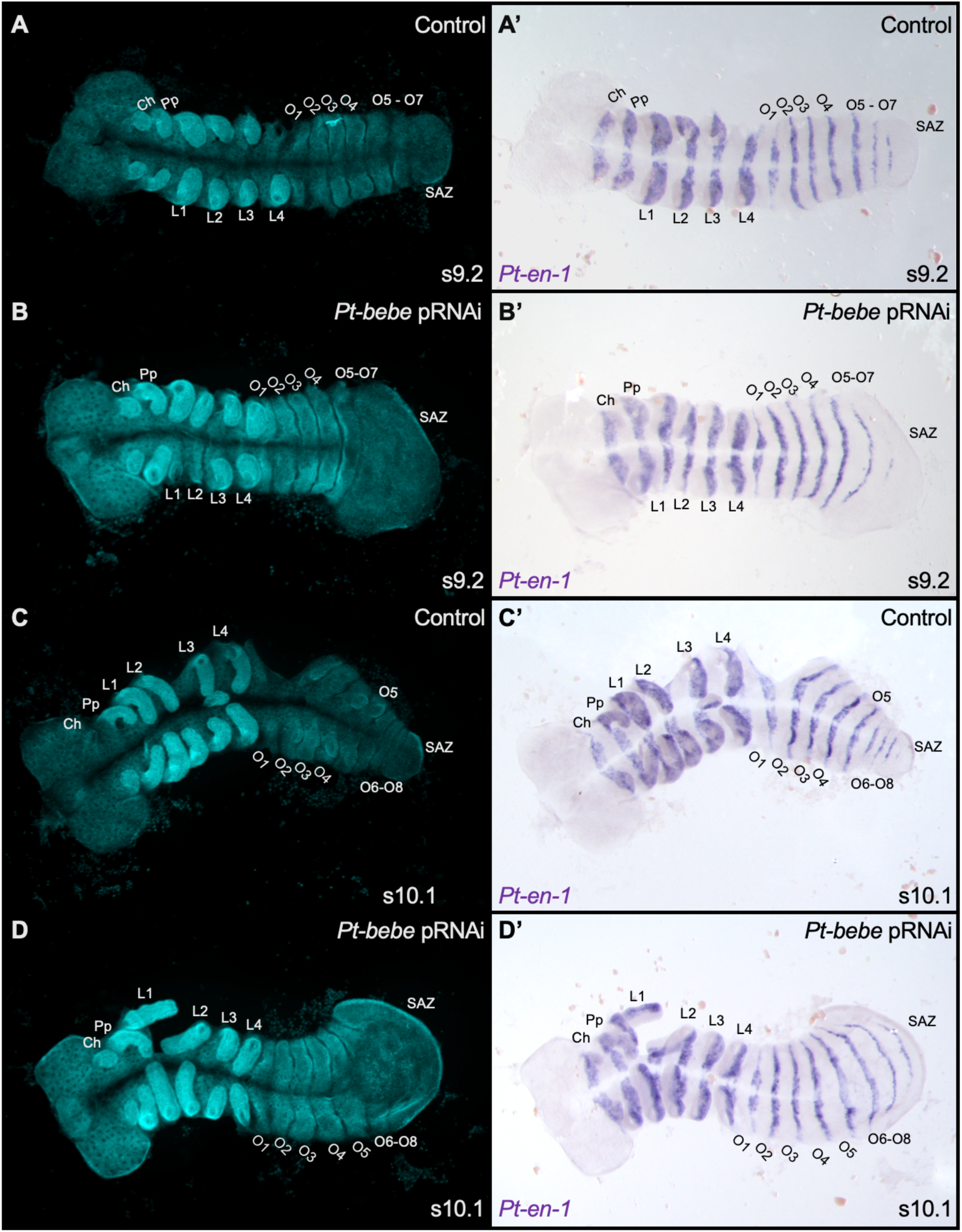
*Pt-en-1* expression in control and *Pt-bebe* pRNAi embryos. Control embryos with DAPI staining (A and C) or under brightfield showing *Pt-en-1* expression (A’ and C’) at stages 9.2 and 10.1. *Pt-bebe* RNAi embryos with DAPI staining (B and D) or under brightfield showing *Pt-en-1* expression (B’ and D’) at stages 9.2 and 10.1. Ch, cheliceral segment; Pp, pedipalpal segment; L, leg-bearing segment; O, opisthosomal segment; SAZ, segment addition zone. Embryos are flat-mounted with the anterior to the left.

Using *Pt-en-1* as a marker, we examined how many cells long (A-P) and wide (D-V) opisthosomal segments were in *Pt-bebe* pRNAi embryos compared to controls (Fig. S18). The A-P length of segments O3-O6 did not differ significantly between *Pt-bebe* pRNAi knockdown and controls (Fig. S18A); nor did the D-V width of segments O2 and O3 (Fig. S18B). However, segments O4 and O5 contained significantly more cells along the D-V axis in *Pt-bebe* pRNAi embryos than controls (Fig. S18B). This is consistent with the increased severity of the *Pt-bebe* pRNAi phenotype in the more posterior opisthosomal segments, i.e., the observed ‘ballooning’ along the D-V axis towards the posterior. To test whether this was due to enlargement of either the dorsal and/or ventral regions of these segments, we assayed the expression of *optomotor blind* (*omb*) and *short gastrulation* (*sog*), which are dorsal and ventral markers, respectively. Expression of both genes was expanded along the D-V axis in the posterior of developing opisthosomas of *Pt-bebe* knockdown embryos, indicating that both the dorsal and ventral regions of segments were enlarged (Fig. S19).

### The SAZ is expanded in *Pt-bebe* knockdown embryos

It was previously shown that expression of *Pt-Wnt8* and *Pt-bebe* overlap in the posterior SAZ^25^. Therefore, we assayed *Pt-Wnt8* expression in *Pt-bebe* pRNAi embryos and *vice versa* to explore their potential interaction. In *Pt-bebe* pRNAi embryos, *Wnt8* expression in the SAZ is broader and extends dorsally around the end of the germband (Fig. 6). This suggests that the SAZ is larger in *Pt-bebe* pRNAi embryos leading to the development of enlarged segments.

**Figure 6.**
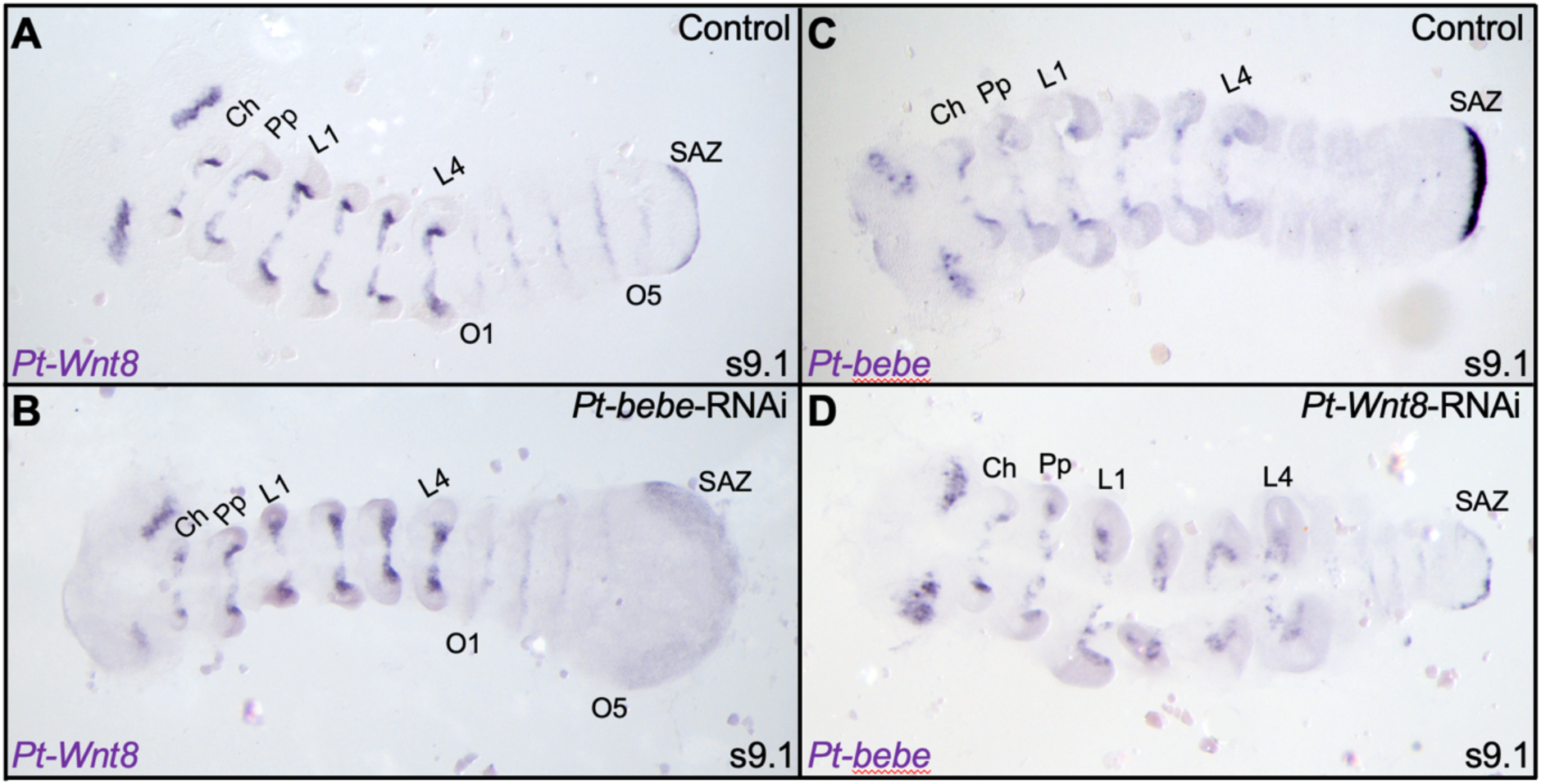
Reciprocal pRNAi knockdown and expression of *Pt-bebe* and *Pt-Wnt8*. Expression of *Pt Wnt8 in* control (A) and *Pt-bebe* RNAi (B) embryos. Expression of *Pt-bebe* in control (C) and *Pt-Wnt8* RNAi (D) embryos. Embryos are flat mounted with anterior to the left. Ch, cheliceral segment; Pp, pedipalpal segment; L, leg-bearing segment; O, opisthosomal segment, SAZ, segment addition zone.

pRNAi knockdown of *Pt-Wnt8* resulted in embryos with narrower opisthosomas consisting of malformed and fewer segments, and one or more smaller SAZs, as observed previously^15^ (Fig. 6). In these embryos we observed weaker, more patchy expression of *Pt-bebe* in the SAZ (note that the other expression domains of *Pt-bebe* appeared to be unaffected) (Fig. 6), although this could also be due to the general reduction and disorganisation of the SAZ in *Pt-Wnt8* knockdown embryos.

These results indicate that *Pt-bebe* and *Pt-Wnt8* interact to regulate the size of the SAZ and consequently the size of segments produced, with *Pt-bebe* constraining the size of the SAZ and opisthosomal segments, since reduced *Pt-bebe* expression results in expansion of the *Wnt8* expression domain and the SAZ.

### *Pt-bebe* may regulate cell organisation/intercalation

The enlargement of the SAZ resulting in wider opisthosomal segments with more cells after *Pt-bebe* pRNAi suggests that this gene may regulate either cell division and/or cell organisation and intercalation of cells. However, there was no obvious increase in the number of dividing cells in the formed segments or SAZ upon *Pt-bebe* pRNAi (Fig. S20).

Cell intercalation during axis elongation in arthropods is regulated by *Toll* genes, encoding a family of cell-surface receptors^30^. In *P. tepidariorum*, knockdown of the *Toll* gene *Pt-LotoA* results in wider segments along the D-V axis but no increase in A-P segment length^30^ – a phenotype strikingly similar to the effect of *Pt-bebe* RNAi knockdown in the opisthosoma (Figs 4 to 6). *Pt-LotoA* is expressed in stripes in all opisthosomal segments in wildtype *P. tepidariorum* embryos and even in the disorganised opisthosoma of *Pt-Wnt8* RNAi embryos^30^ (Fig. S21). However, in *Pt-bebe* pRNAi embryos, *Pt-LotoA* expression appears weaker in posterior opisthosomal segments relative to anterior ones, reflecting the increased severity of the *Pt-bebe* knockdown phenotype from anterior to posterior (Fig. S21). This suggests that *Pt-bebe* knockdown may perturb cell organisation in the opisthosoma by potentially inhibiting *Pt-LotoA* expression, at least indirectly.

## Discussion

### Phylogenetic distribution and potential loss of *bebe* during evolution

Our study shows that *bebe* is a newly identified but ancient animal toolkit gene. However, *bebe* has a rather unusual phylogenetic distribution. We could not identify *bebe* orthologues in either mammals or polyneopteran and holometabolous insects, which may help explain why this gene has been overlooked to date. *bebe* also appears to have been lost in specific lineages within phyla. For example, although *bebe* is found in hemimetabolous insects, arachnids, annelids and nematodes, we could not find orthologues in dragonflies, harvestmen, the annelid *P. dumerilii*, or the nematode *Caenorhabditis elegans*, as well as other species in these phyla (Supplementary File 1). This suggests that despite the ancient origin and likely important roles of *bebe* in the development of many animals, it can also be readily lost during evolution.

Comparative genomics has shown that gene loss is common in evolution^31^, including other toolkit genes, for example, the loss of *Wnt2* in insects and *Wnt3* in protostomes^29^. In the case of *bebe,* because this gene does not appear to be part of a multi-gene family or even closely related to other genes, its loss is even more unusual. We speculate that it may have been lost through functional redundancy or that other genes replaced its role in lineages where it is not found, although these replacements must belong to distinct gene families, albeit that potentially play similar roles and interact with the same proteins as Bebe. Further understanding of this process requires more knowledge of *bebe’s* role and the proteins it interacts with in a range of animals.

### Bebe regulates SAZ and segment size in *P. tepidariorum* potentially through interactions with Wnt signalling

We demonstrate that *bebe* regulates head lobe development, and SAZ formation and segment size in the spider *P. tepidariorum.* The expression of *bebe* orthologues suggests that where retained, it likely also plays similar roles in other segmented animals, except for vertebrates. Knockdown of *bebe* expression in *P. tepidariorum* resulted in enlargement of the SAZ and wider posterior segments containing more cells, although most opisthosomal segments were still produced. This suggests that segment size can be specified by the SAZ but that this can be decoupled from the mechanisms that sequentially generate segments from this tissue.

Given that in wildtype embryos, the *bebe* expression domain reduces in size concomitantly with the SAZ as segments are formed, and that knockdown of *bebe* increases the size of the SAZ, we conclude that *bebe* acts to limit SAZ size in *P. tepidariorum*. Conversely, *Wnt8* promotes SAZ size because knockdown of *Pt-Wnt8* reduces the SAZ^15^. The overlap in *Pt-bebe* and *Pt-Wnt8* expression, the dependence of the posterior *Pt-bebe* expression domain on *Pt-Wnt8* and the opposite effects of the *Pt-bebe* and *Pt-Wnt8* knockdowns suggest that these two genes are parts of the same gene regulatory network specifying the size of the SAZ, and consequently segments. In this network, *Pt-bebe* and *Pt-Wnt8* appear to play opposing roles. As *Pt-bebe* expression is reduced upon *Pt-Wnt8* pRNAi, we speculate that the latter may activate the former in the SAZ. It is possible that the level of *Pt-bebe* expression in the SAZ is also regulated by Hh because *bebe* expression was shown to be negatively regulated by Hh in a previous transcriptomics screen for Hh targets^27^. We suggest that Pt-Bebe then in turn feeds-back to modulate Wnt signalling. Given that *bebe* is predicted to encode a secreted protein it could potentially regulate Wnt signalling outside the cell through direct modulation of the activity Wnt8 or other Wnt ligands expressed in the SAZ of spiders and other panarthropods^29,32-35^. Therefore, Bebe could play a similar role to the glypicans Dally and Dally-like, which can sequester Wnt ligands and modulate Wnt gradients^36,37^. Alternatively, it could even act as a ligand itself through a known or an unknown receptor.

We also found evidence that *Pt-bebe* may regulate *Pt-LotoA* during segmentation in *P. tepidariorum*. Knockdown of *Pt-LotoA* produces wider segments very similar to the *Pt-bebe* phenotype^30^. *Pt-LotoA* expression does not overlap with *Pt-bebe*, but the related gene *Pt-LotoB* is expressed in the SAZ (although knockdown of this gene alone does not affect segment size)^30^. It is therefore possible that the D-V widening of segments seen in *Pt-bebe* pRNAi embryos results from disrupted cell organisation via perturbation of Toll signalling.

### Other roles of *bebe* in animal development

In early *P. tepidariorum* embryos, *Pt-bebe* is expressed around the blastopore (this work,^27^) where it likely helps specify the SAZ before segmentation begins, as discussed above. Intriguingly, *bebe* is also expressed around the blastopore in nearly all the other species we surveyed. For example, early expression of *bebe* is similar to that of *brachyury* in the millipede *G. marginata* and the onychophoran *E. kanangrensis*^38^, and *bebe* is expressed in potential mesoderm around the blastopore of the annelid *O. fusiformis* as well as in the blastopore lip of the cnidarian *N. vectensis*. Across metazoans, including *N. vectensis*, the blastopore region is a centre of high Wnt/ß-catenin signalling activity^39^. Interestingly, analysis of the previously published RNA-seq dataset^40^ shows that *bebe* is strongly upregulated in *N. vectensis* embryos with pharmacologically or genetically upregulated β-catenin signalling (8.65x in gastrulae treated with GSK3β inhibitor 1-azakenpaullone (AZK) since early blastula in comparison to DMSO controls; 19.6x in 3 day larvae treated with AZK since early blastula in comparison to DMSO controls, and 59.97x in 3 day APC^-/-^ mutant larvae in comparison to wild type). Our data therefore suggest that *bebe* could play an early role in modulating Wnt activity in this region of the embryos of animals where it has been retained, and perhaps even ancestrally. However, since Wnt signalling is not required for the morphogenetic movements of gastrulation in *N. vectensis*^41^, it would point more to an early role for *bebe* in Wnt-dependent axial patterning, which is a highly conserved feature across Metazoa.

*bebe* expression in *D. rerio* may be derived compared with the other animals, as it likely does not play a role in axial patterning or segmentation in this species, although it is expressed in tissues that also express Wnts in this vertebrate^42^. The earliest expression we detected was at 16 hpf in presumptive placodal cells and then later in potentially in cranial neural crest cells and subsequently in the pharyngeal arches. However, the neural expression may be conserved, as *bebe* orthologues also appear to be expressed in developing brain regions of most bilaterian species we surveyed. Consistent with this, knocking down *bebe* in the spider *P. tepidariorum* results in disorganised head lobes. Our functional analysis of *bebe* in the spider, together with its expression in the head and around the blastopore of other animals, suggests it may play a general role in modulating Wnt signalling in these developing tissues.

### Conclusions

We have identified *bebe* as a novel but ancient toolkit gene that regulates SAZ and segment size in a spider and likely plays important roles in the development of other animals. However, *bebe* has a striking phylogenetic distribution and appears to have been lost in several lineages, including mammals and major insect groups. Our study emphasises the need for unbiased approaches in a wider range of animals to understand the regulation and mechanisms of development. Future analyses of *bebe* function in additional animals are needed to reveal more about the role and interactions of this ancient gene and may point to why it has become dispensable in some animals during evolution.

## Materials and Methods

### Parasteatoda tepidariorum

*P. tepidariorum* adults were obtained from the Durham and Munich laboratory colonies, and embryos were staged and fixed as described previously^43,44^. Stages 6-10.2 were selected as the SAZ is set up at around stage 6 where the first segment (O1) appears, followed by sequentially forming segments in stages 7 to 10.1.

RNA was extracted from embryos stages 5-10.2 using TRIzol™ Reagent (Thermo Fisher Scientific). cDNA was synthesised using the High-Capacity cDNA Reverse Transcription Kit (Applied Biosystems™). The *Pt-bebe* sequence was obtained from the *P. tepidariorum* transcriptome^25^. A 710 bp fragment of the *Pt-bebe* coding region was amplified using primers designed with Primer3 (http://primer3.ut.ee) (Supplementary File 1).

RNA probes were labelled for either colorimetric or fluorescent *in situ* hybridization (FISH) as previously described with minor modifications^45^. Nuclear staining was performed by incubating embryos in 1 mg/ml 4-6 diamindino-2-phenylindol (DAPI) in PBS with 0.1% Tween-20 for 30 min. Segment identity in stained embryos was determined by morphological markers in images of DAPI staining.

Fragments of the coding region of *Pt-bebe* (710 bp) and GFP were cloned using a TOPO TA cloning kit (Thermo Fisher Scientific) and amplified using T7 and T3/T7 adaptor primers (Supplementary File 1). *In vitro* transcription of both strands was then carried out using the MegaScript T7 transcription kit (Invitrogen) to generate double stranded RNA (dsRNA). The concentration was then adjusted to 1.5-2.0 ng/µl and stored at -20°C.

*Pt-en-1*, *Pt-Wnt8*, *Pt-sog*, *Pt-omb*, *Pt-eve* sequences for RNA probes and/or doubles stranded RNA syntheses were amplified as described previously^15,24,46^.

Adult female spiders were injected according to the protocol by Akiyama-Oda and Oda^46^. 1.5-2 µl of dsRNA was injected into the opisthosoma at concentrations of 1.5-2.0 ng/µl every two to three days for a total of five injections per animal. At least four control females were injected with double stranded green fluorescent protein (GFP) as above. RNAi and control females were mated after the second injection. By the 3^rd^ cocoon laid after mating, embryos from all injected spiders were collected for gene expression and phenotypic analyses.

For immunohistochemistry, spider embryos were fixed (1 ml 2x PBS or (2x PEMS), 1 ml 4% PFA) and about 2 ml heptane) for 3 to 5 hours, followed by washes in MEOH, and then kept at - 20°C until use. Fixed embryos were gradually washed from 100% MEOH in 1% Triton-X in 1X-PBS (1% PBT). Subsequent washes in 1% Triton-X in 1X-PBS followed until embryos were permeabilised for 15 min with 3% Triton-X in 1X-PBS (3% PBT). Embryos were blocked in Blocking in 5% normal goat serum (NGS) for about 1 hr at RT. The primary antibody, Phospho-Histone H3 (Ser10) Antibody (#9701 from Cell Technology) was added at a 1:1000 dilution and left overnight at 4°C. Embryos were washed with 1% PBT several times to ensure no trace of the primary remained, and the primary antibody was recovered and stored at 4°C for future use.

To count cells in embryos, fluorescent in situ hybridisation was performed to detect segment boundaries marked by *Pt-en-1* in wild type and *Pt-bebe* pRNAi embryos. Cells marked by *en-1* staining were manually counted in ImageJ/Fiji^47^ along the D-V axis (total length of the stripe) for the opisthosomal segments (O2, O3, O4, and O5). For the same opisthosomal segments, cells were counted along the A-P axis (the width). Five embryos were analysed for the control and *Pt-bebe* pRNAi embryos. A Mann-Whitney U test (non-parametric) was used to test for a significant difference between control embryo cell count and *bebe* pRNAi embryos. This test was conducted for each stripe length and segment width.

### Acanthoscurria geniculata

A 1624 bp fragment of *A. geniculata bebe* spanning the complete CDS of the gene, was amplified using specific primers (Supplementary File 1). A 1141 bp probe template was generated via semi-nested PCR with an internal reverse primer (Supplementary File 1). An anti-sense DIG labelled RNA probe (DIG RNA Labelling Mix; Roche) of *A. geniculata bebe* was generated using T7 RNA polymerase (Roche).

Staging of *A. geniculata* embryos was carried out according to Pechmann (2020)^48^. In situ hybridisation in *A. geniculata* was performed as described in Pechmann (2020)^48^. In situ stained embryos were incubated for one hour in a PBST/Sytox green solution (final concentration of Sytox green: 1µmol) and images were recorded using an AxioZoom.V16 (Zeiss; ZEN2 Software) equipped with an AxioCam 506 colour camera. Projections of image stacks were carried out using Helicon Focus (HeliconSoft) and overlays were created using Photoshop CS5.1.

### Pardosa amentata, Pholcus phalangioides, Glomeris marginata, Pyrrhocoris apterus, and Euperipatoides kanangrensis

For all species, fragments of the *bebe* orthologous were isolated by means of RT-PCR using mRNA isolated (Dynabeads mRNA Purification Kit (Invitrogen)) from embryonic total RNA extractions (TRIZOL (Invitrogen)). Gene specific primers were used that we retrieved from embryonic transcriptomes (Supplementary File 1). In all cases a nested primers were used to boost the amplification and to add higher specificity (Supplementary File 1). All nested backward/reverse (bw) primers were equipped with a 5′-T7 promotor site for probe synthesis^49^.

For all species, the same whole-mount in-situ hybridization (WISH) protocol was used^50^. After WISH, embryos were incubated in 1:10,000 SYBR-green or 1:1000 DAPI in PBST with 0.1% Tween-20 for approximately 30 minutes. Unincorporated dye was removed by repeated washing steps with PBST. Pictures were taken under a MZ-FLIII Leica dissection microscope with a Leica DC490 digital camera and an external UV lamp. Linear adjustments were made using the image-processing software Adobe Photoshop 2022.

### Owenia fusiformis

Gravid adult *O. fusiformis* were collected from Roscoff, France. Adult worms were spawned via *in vitro* fertilisation, the resulting larvae reared and fixed by established methods^51,52^

Single colorimetric whole-mount *in situ* hybridisation was performed as in Carrillo-Baltodano et al^52^. A 1308 bp fragment of *Of-bebe* was amplified using specific primers to generate a template for riboprobe synthesis (Supplementary File 1). In brief, methanol stored animals were rehydrated in PBS+0.1% Tween-20 (PTw), permeabilised with 5 μg/ml Proteinase-K for 90 s (ThermoFisher Scientific), and hybridised with 1 ng/µl of RNA probe for 48 h at 62 °C, before colourimetric development steps. Stained animals were mounted in glycerol and imaged.

Differential interface contrast (DIC) imaging was performed using a Leica CTRMIC compound microscope with a mounted Luminera Infinity5 camera. Images were digitally stacked using Helicon Focus 8 (HeliconSoft), and further editing of brightness/contrast was performed using Adobe Photoshop (Adobe Systems, Inc.).

### Danio rerio

Wild-type WIK *Danio rerio* were maintained at 28°C on a 14h light/10h dark cycle. Zebrafish experimental procedures and husbandry were conducted under personal and project licences granted by the UK Home Office under the United Kingdom Animals and Scientific Procedures Act (ASPA) and following ethical policies approved by the University of Exeter’s Animal Welfare and Ethical Review Body (AWERB). Experiments on zebrafish embryos were carried out before animals became capable of independent feeding at 120 hpf.

A partial *bebe* sequence was amplified from zebrafish cDNA generated from RNA isolated from 1 dpf pooled zebrafish embryos using gene specific primers (Supplementary File 1).

The PCR fragment was ligated into the pCRII-TOPO TA vector using the TOPO TA cloning Kit (Invitrogen). The antisense *Dd-bebe* digoxigenin probe was transcribed using SP6 RNA Polymerase (Promega) and DIG RNA labelling detection kit (Roche). The DIG probe was purified through MicroSpinTM G-50 columns (GE Healthcare).

Wild-type embryos were collected and fixed at 16, 20, 24, 30 and 48hpf with 4% PFA overnight. *In situ* hybridisation was carried out as previously described^53^. Stained embryos were kept in 70% glycerol and imaged with an Olympus SZX16 stereo microscope equipped with a DP71 digital camera. Images were taken using the Olympus cellSens Standard software. For flat-mounted dorsal views, the yolks were removed with forceps and embryos mounted on a slide between two coverslips ventral side down.

### Nematostella vectensis

A fragment of the *N. vectensis* ortholog of *bebe* was isolated from mixed embryonic stages cDNA by PCR (Supplementary File 1) and cloned in pJet1.2 (Thermo Fisher). Dig-labelled RNA probe was generated with a HighScribe T7 kit (NEB). For in situ hybridization in *N. vectensis*, embryos were fixed in 4% PFA/PBS for 1 hour at room temperature, washed three times with PTw (1x PBS, 0.1% Tween20), then twice in 100% methanol, and stored in 100% methanol at -20°C. To avoid contraction upon fixation, primary polyps were relaxed prior to fixation by addition of 1-2 ml of 1M MgCl_2_ per 10 cm Petri dish with polyps in *N. vectensis* medium. The embryos were rehydrated via a single 50% methanol/50% PTw and three PTw steps, and digested with 10 µg/ml proteinase K/PTw for 10 minutes. The digest was stopped by two sequential washes in 4 mg/ml glycine/PTw, the embryos were washed twice in 1% triethanolamine/PTw (TEA), then once with 3 µl/ml acetic anhydride/TEA, and once in 6 µl/ml acetic anhydride/TEA. Then, the embryos were washed twice with PTw and post-fixed for 1 hour in 3.7% formaldehyde/PTw. Formaldehyde was removed by five PTw washes, and the embryos were transferred into hybridization solution (Hybe: 50% formamide, 5x SSC, 1% SDS, 0.1% Tween20, 5 mg/ml torula yeast RNA, 50 µg/ml heparin) in two 10-minute steps: first, 50% Hybe/50% PTw, then 100% Hybe. Then, the embryos were prehybridized in Hybe for 2 hours at 60°C and hybridized overnight with 0.5 ng/µl *bebe* probe diluted in Hybe. Unbound probe was removed by a series of 30-minute washes at 60°C : first in 100% Hybe, then in 75% Hybe/25% 2xSSCT (2x SSC, 0.1% Tween20), then in 50% Hybe/50% 2x SSCT, then in 25% Hybe/75% SSCT, then in 2x SSCT. After this wash, 4d planulae and primary polyps were incubated for 30 minutes in 1 U/µl RNase T1/2x SSC at 37°C to reduce unspecific pharyngeal staining. Subsequently, the samples were washed thrice for 20 minutes in 0.2x SSCT at 60°C, and moved to PTw at room temperature. After 1 hour blocking in 1% Roche blocking reagent/MAB (Block), the embryos were stained overnight at 4°C in anti-Dig-AP Fab fragments (Roche) diluted 1:2000 in Block. Unbound antibody was removed by ten 10-minute washes in PTw, the embryos were washed twice with AP buffer and stained with 2 µl/ml of NBT and 2 µl/ml BCIP (both Roche) diluted in the AP buffer. After staining, the embryos were rinsed once with water and left in ethanol for several hours. Then, after a short PTw rinse, the embryos were infiltrated with 86% glycerol and imaged using under a Nikon Eclipse 80i microscope equipped with a Nikon DS-Fi3 camera.

### Phylogeny and amino acid alignment

We surveyed available genomic and transcriptomic resources on GenBank for orthologues of *bebe*, as well as arachnid transcriptomes^54^ using BLAST searches^55^. We aligned sequences with MAFFT using the einsi setting and the gap extension parameter set to 0^56^, and trimmed the alignment with TRIMAL with a similarity threshold of 0.3, gap threshold of 0.5, and consistency threshold of 20^57^. We inferred a maximum likelihood phylogeny with IQ-TREE version 3.0.1 using ModelFinder to choose the substitution model (Q.PFAM+I+R6) and 10,000 ultrafast bootstrap replicates to determine branch support^58-60^. The tree was visualized in FigTree version 1.4.4 [https://tree.bio.ed.ac.uk/software/figtree/]. To characterize the possible features and functions of this undescribed protein we searched for its orthologues in UniProt and used InterProScan, Phobius, SignalP6.0 and AlphaFold to annotate domains^61-64^.

## Data availability statement

All data and resources associated with this work are available in the manuscript or supplementary material. New sequence resources for *P. phalangioides, G. marginata, P. apterus,* and *E. kanangrensis* have been deposited in FigShare: https://doi.org/10.6084/m9.figshare.34021908.

## Acknowledgements

This study was funded by a studentship from Durham University to ZXS, a Nigel Groome studentship from Oxford Brookes University to GB and grants from the Leverhulme Trust (RPG-2016-234) and the NERC (NE/T006854/1) to APM. The work of RJ was funded by the Swedish Natural Science Council (VR) (grant no. 2022–03522). Work in the Genikhovich group is funded by the Austrian Science Fund (FWF) grants PAT 2395824 and P36080 to GG. The work in the Turetzek and Pechmann group was funded by the Deutsche Forschungsgemeinschaft (DFG, German Research Foundation)—project number 503325252, part of the Priority Programme SPP 2349: Genomic Basis of Evolutionary Innovations (GEvol, project number 458908270). CMD and TF were supported by a Leverhulme Trust grant (RPG-2023-121). The work of GS and SS was funded by the BBSRC Research Grant UKRI725. We thank Saul Mallinson for advice on data analysis.

