## Supplementary figures for "Lost and found: *beer belly* is a novel, ancient regulator of animal development"

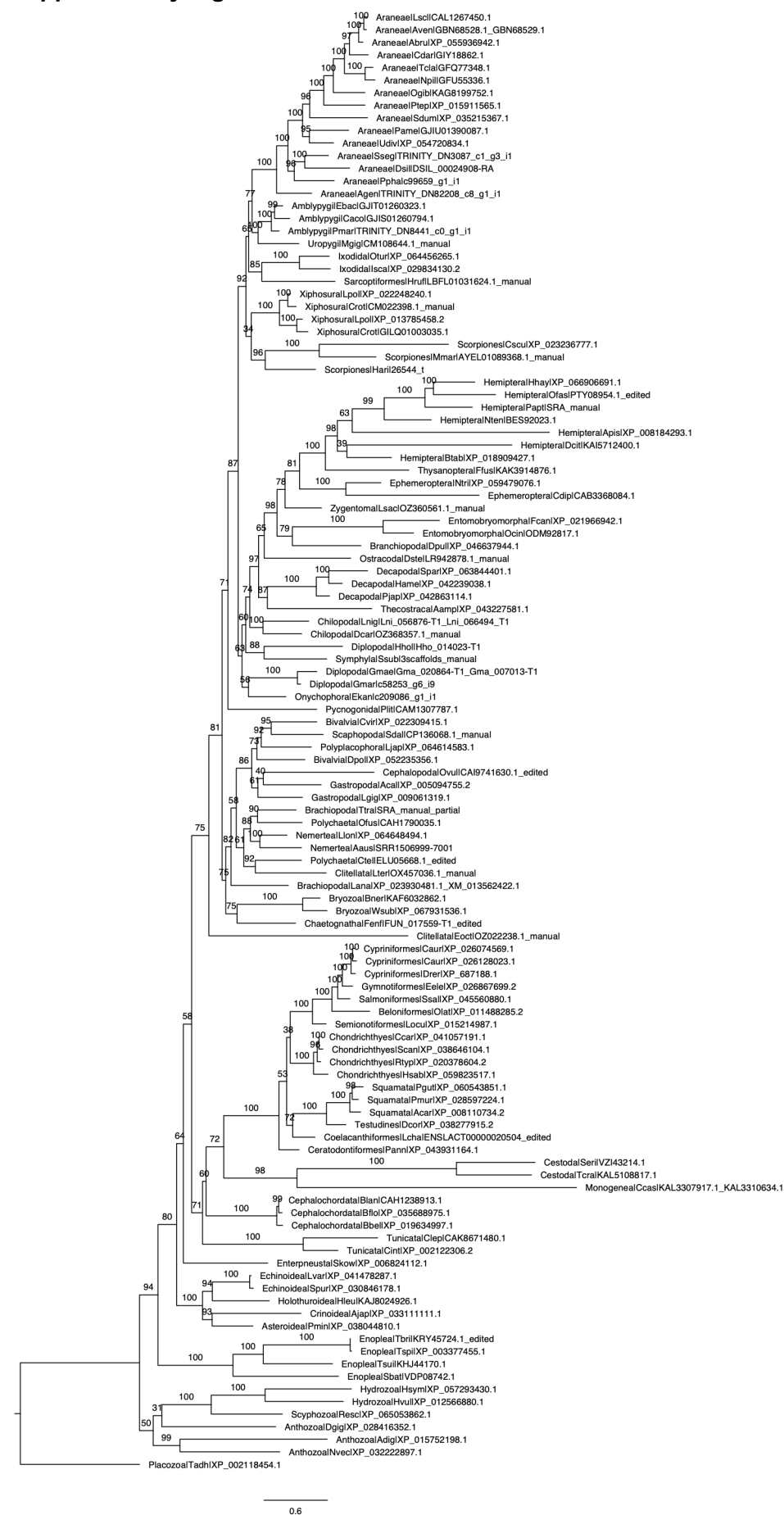

**Figure S1. Phylogeny of Bebe orthologues.** Maximum likelihood phylogeny of Bebe orthologues inferred using with IQ-TREE (version 3.0.1) using ModelFinder to choose the substitution model (Q.PFAM+I+R6) and 10,000 ultrafast bootstrap. See Supplementary File 1 for sequences.

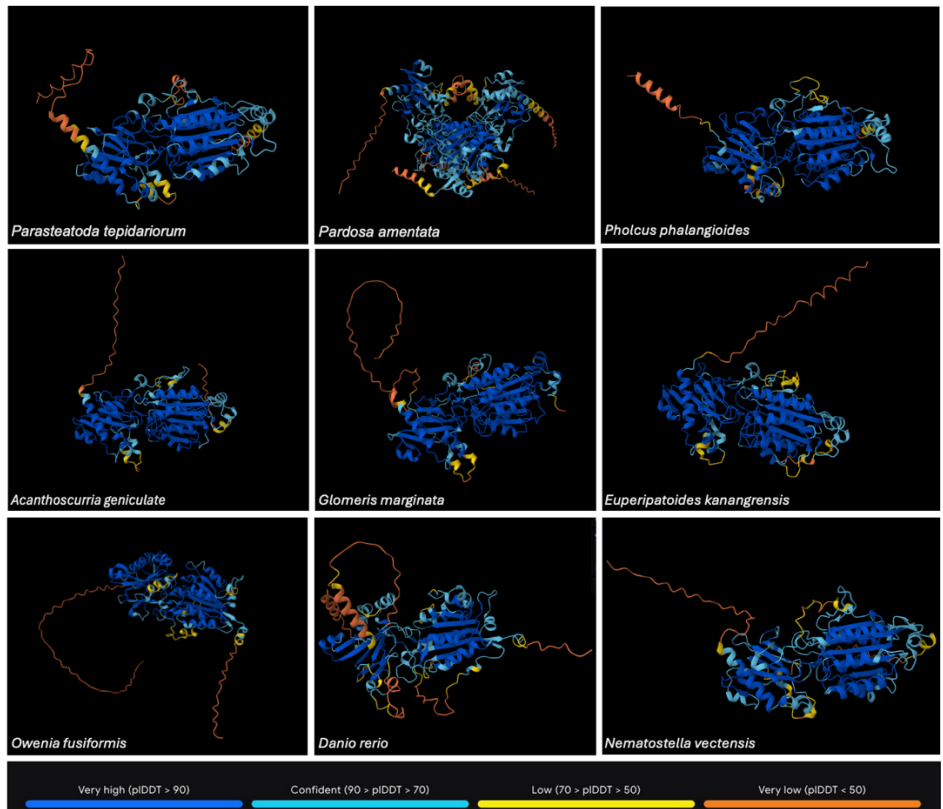

**Figure S2. Predicted structure of Bebe in *P. tepidariorum* and other selected species.** AlphaFold predictions for the Bebe orthologues of the spiders *P. tepidariorum*, *Pholcus phalangioides*, and *Acanthoscurria geniculata*, the millipede *Glomeris marginata*, the hemipteran insect *Pyrrhocoris apterus*, the onychophoran *Euperipatoides kanangrensis*, the annelid *Owenia fusiformis*, the zebrafish *Danio rerio*, and the cnidarian *Nematostella vectensis*. Default AlphaFold parameters were used.

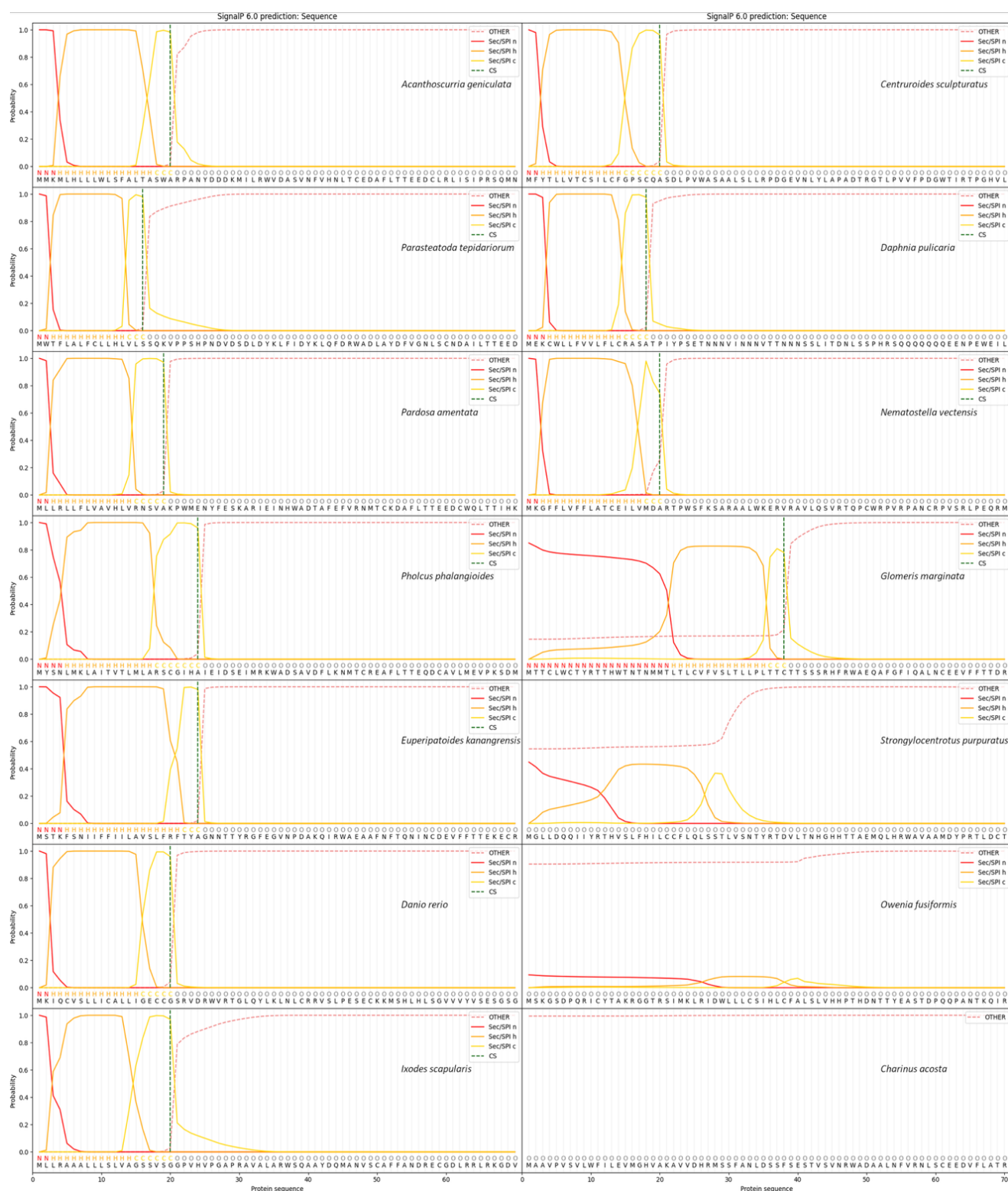

**Figure S3. Prediction of Signal Peptides in the Bebe orthologues.** Presence of an N-terminus signal peptide was predicted using SignalP6.0 across selected species except for *O. fusiformis*. The prediction indicates the probability of a signal peptide at a specific residue or region of the protein sequence. The type of predicted signal peptide is characterised as follows: Other (red dashed-line; probability of no signal peptide); Sec/SPI n (solid red-line; N-region); Sec/SPI h (solid orange-line; Hydrophobic region); Sec/SPI c (yellow-solid-line; C-region); Cs (green dashed-line; predicted cleavage site).

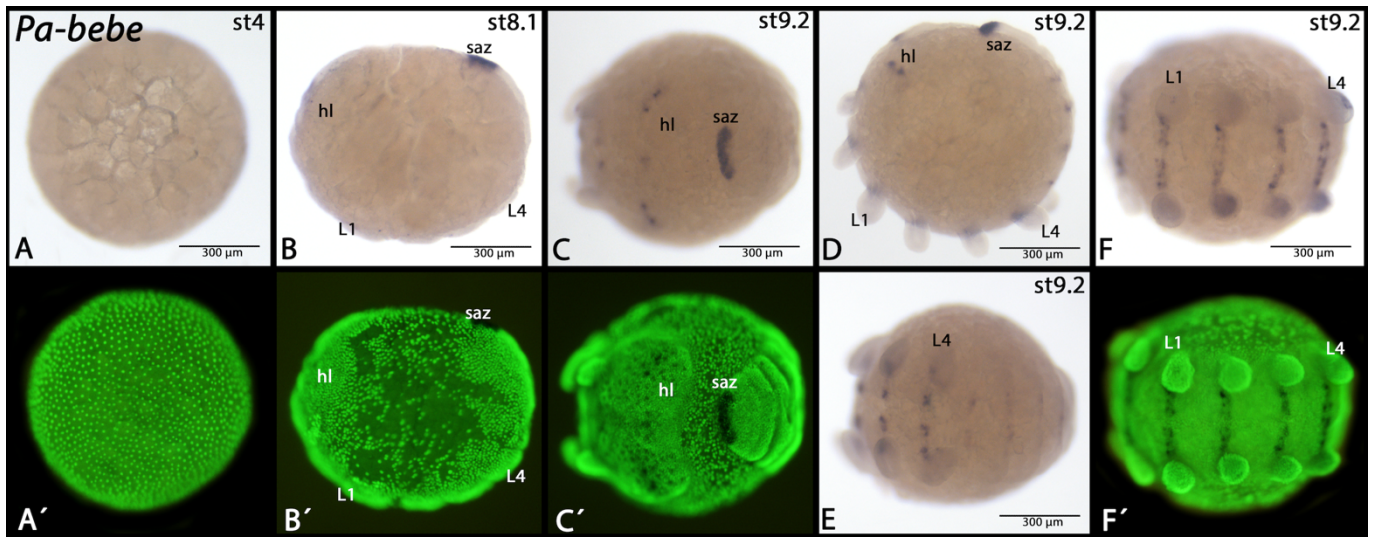

**Figure S4. Expression of *Pardosa amentata bebe*.** In all panels anterior is to the left (except panel B showing a blastoderm stage with unclear orientation). Panels B and D represent lateral views. Panel C shows a view of the head lobe and the posterior SAZ (note that the embryo is curled around the yolk (central mass)). Panels E and F represent ventral views. Panels marked with an apostrophe represent SYBR-green staining of corresponding embryos. Developmental stages are indicated in the bright field panels (staging after<sup>1</sup>). Abbreviations: hl, head lobe; L1/4, first respectively fourth walking leg; saz, segment addition zone.

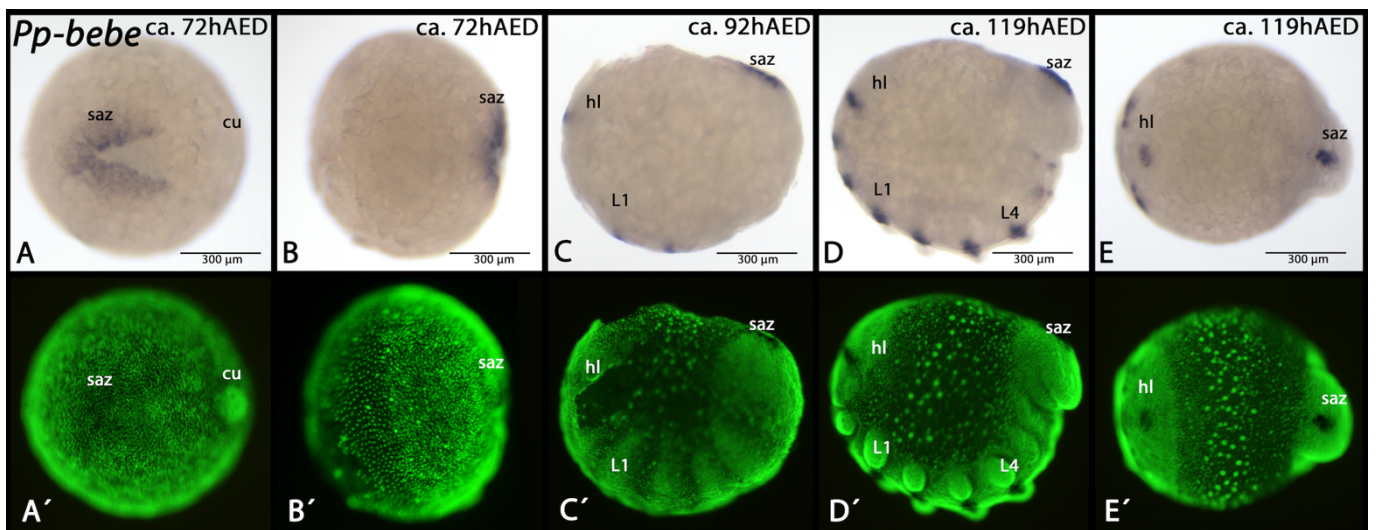

**Figure S5. Expression of *Pholcus phalangioides bebe*.** In all panels, anterior is to the left. Panel A represents a ventral view. Panels B-D represent lateral views. Panel E represents a dorsal view. Panels marked with an apostrophe represent SYBR-green staining of corresponding embryos. Developmental stages are indicated in the bright field panels (staging after<sup>2</sup>). Abbreviations: cu, cumulus; hl, head lobe; L1/4, first respectively fourth walking leg; saz, segment addition zone. Note the image in panel D is also displayed in Fig. 3A.

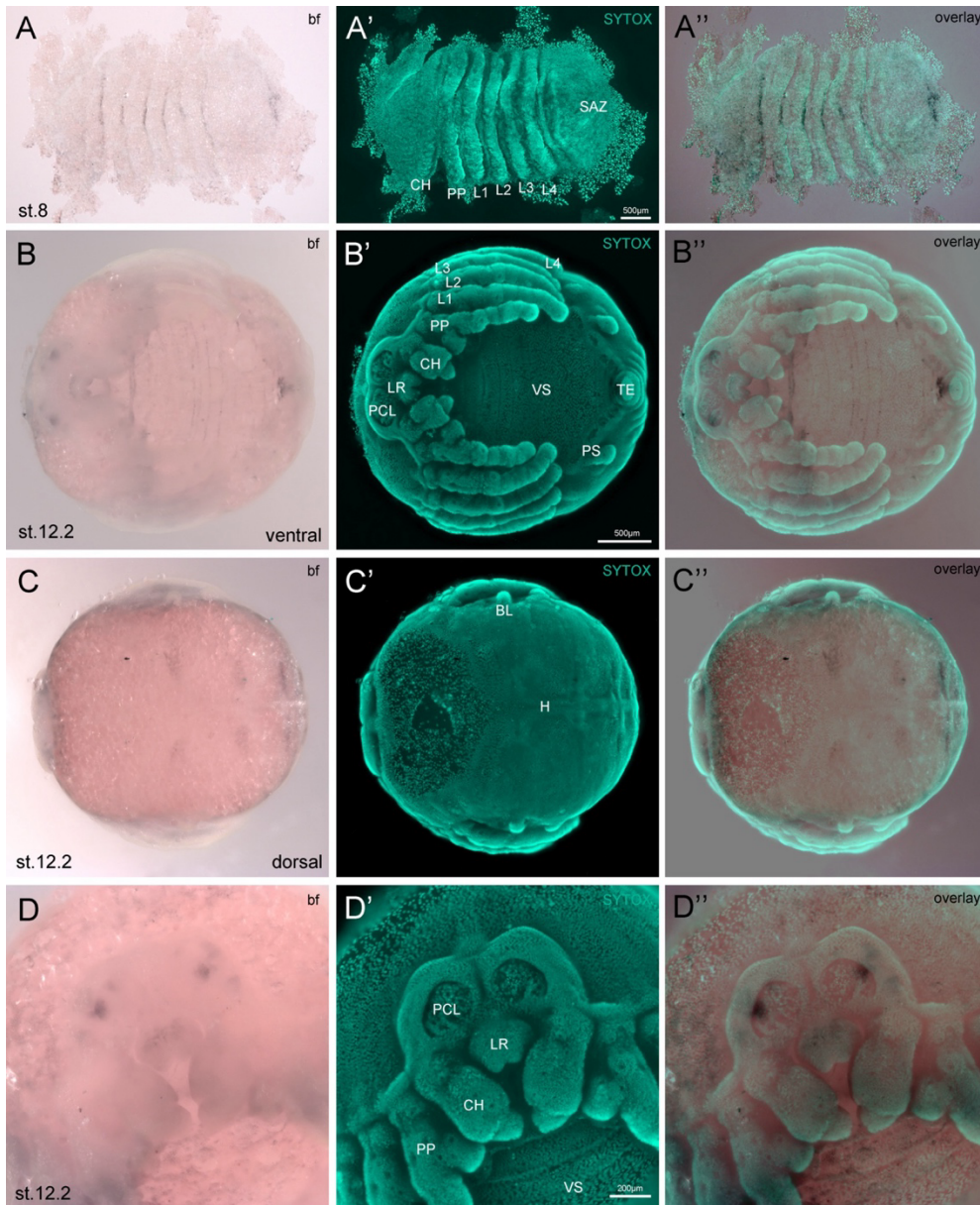

**Figure S6. Expression of *Acanthoscurria geniculata bebe*.** (A) At stage 8, *Ag-beer-belly* is expressed in stripes in the prosomal segments of the walking legs, pedipalps and chelicers. (B-D) At stage 12, *Ag-beer-belly* is expressed in thin transversal stripes within the ventral sulcus and close to the telson (B). Additional expression is visible in spots within the pre-cheliceral lobe, at the base of the labrum (B and D) and in dorsal segmental blocks, left and right to the heart tube (C). Abbreviations: BL - book lung; CH - chelicera; H - heart; L1-L4 - walking legs; LR - labrum; PCL - pre-cheliceral lobe; PP - pedipalp; PS - posterior spinneret; SAZ - segment addition zone; TE - telson; VS - ventral sulcus.

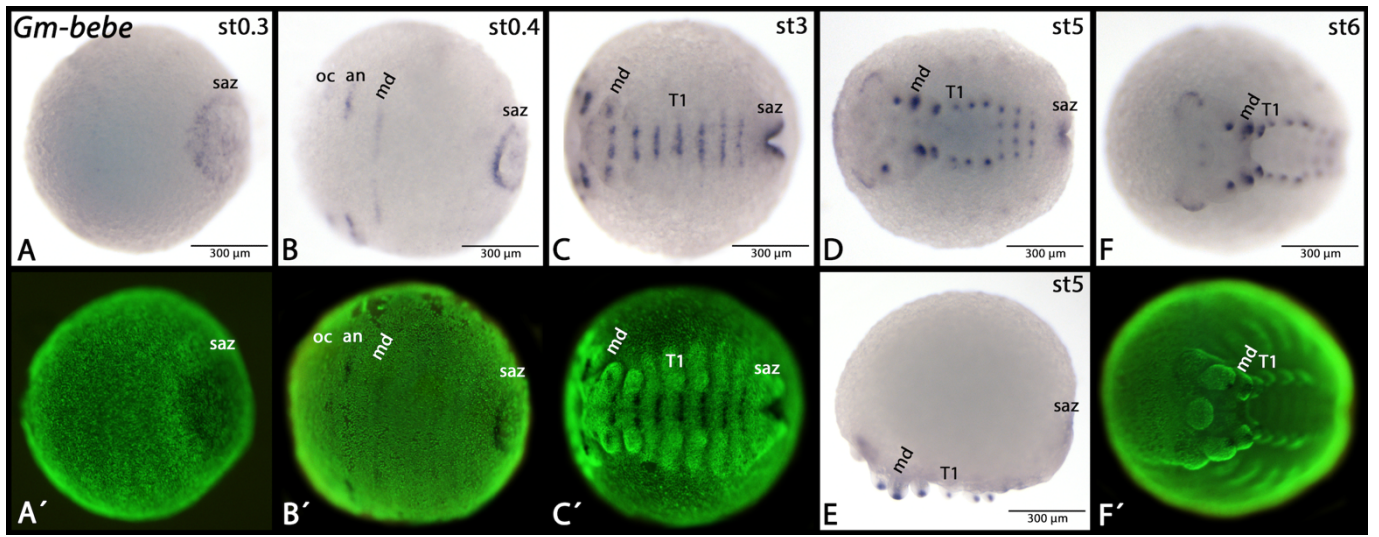

**Figure S7. Expression of *Glomeris marginata bebe*.** In all panels, anterior is to the left. All panels represent ventral views, except for panel E (lateral view). Panels marked with an apostrophe represent SYBR-green staining of corresponding embryos. Developmental stages are indicated in the bright field panels (staging after<sup>3</sup>). Abbreviations: an, antennal segment; md, mandibular segment; oc, ocular region; saz, segment addition zone; T1, first trunk segment. Note the image in panel C is also displayed in Fig. 3C.

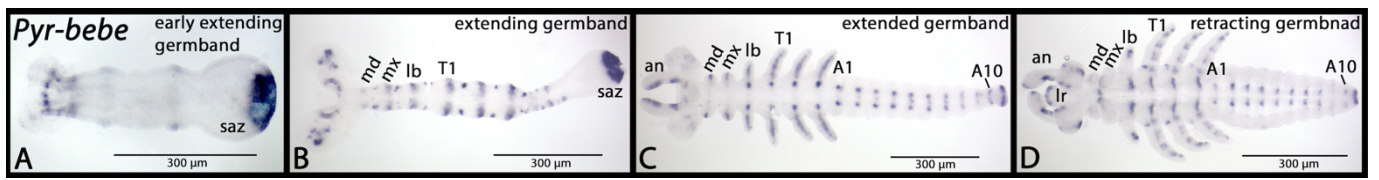

**Figure S8. Expression of *Pyrrhocoris apterus bebe*.** In all panels, anterior is to the left. All panels represent ventral views on flat-mounted embryos. Abbreviations: A1/10, first respectively tenth abdominal segment; an, antennal segment; lb, labial segment; lr, labrum; md, mandibular segment; mx, maxillary segment; saz, segment addition zone; T1, first trunk segment. Note the image in panel B is also displayed in Fig. 3D.

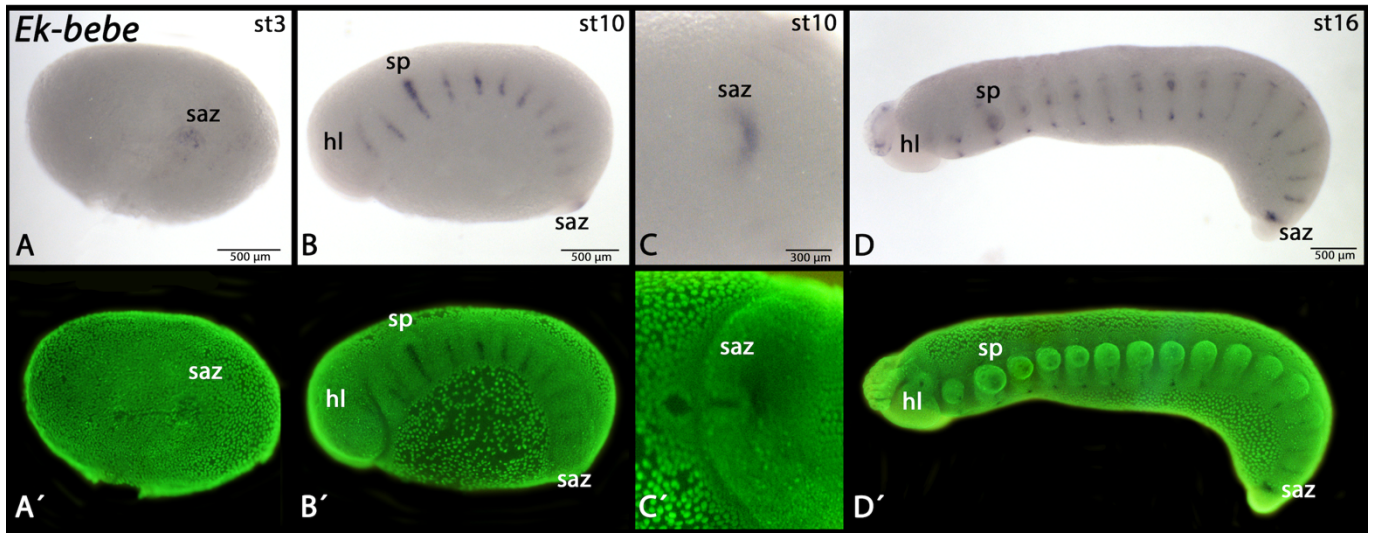

**Figure S9. Expression of *Euperipatoides kanangrensis bebe*.** In all panels, anterior is to the left. Panels A and C represent ventral views, panels B and D represent lateral views. Panels marked with an apostrophe represent SYBR-green staining of corresponding embryos. Developmental stages are indicated in the bright field panels (staging after<sup>4</sup>). Abbreviations: hl, head lobe; saz, segment addition zone; sp, slime papilla. Note the image in panel B is also displayed in Fig. 3E.

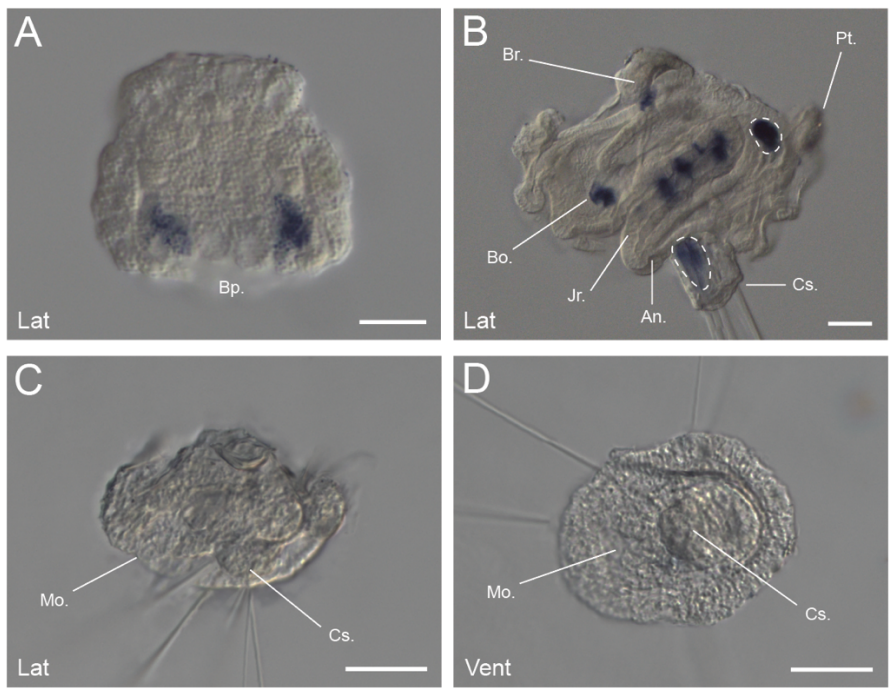

**Figure S10. Expression of *Owenia fusiformis bebe*.** In the gastrula stage of *O. fusiformis*, *bebe* is expressed in the endomesoderm (A). In competent larvae (~21dpf), *bebe* is expressed in paired lateral stripes at the boundaries of developing segments in the juvenile trunk rudiment. Expression is also detected in the buccal organ and the brain. Dashed lines indicate background signal from probe capture in the chaeta-sac and midgut. (B). In the early *O. fusiformis* 'head-larva', prior to the onset of trunk rudiment development (24 hpf), *bebe* is not expressed (C and D). Scale bars = 50 μm. Bp = blastopore; Cs = chaeta-sac; Br = brain; Pt = prototroch; Bo = buccal organ; Jr = juvenile rudiment; An = anus.

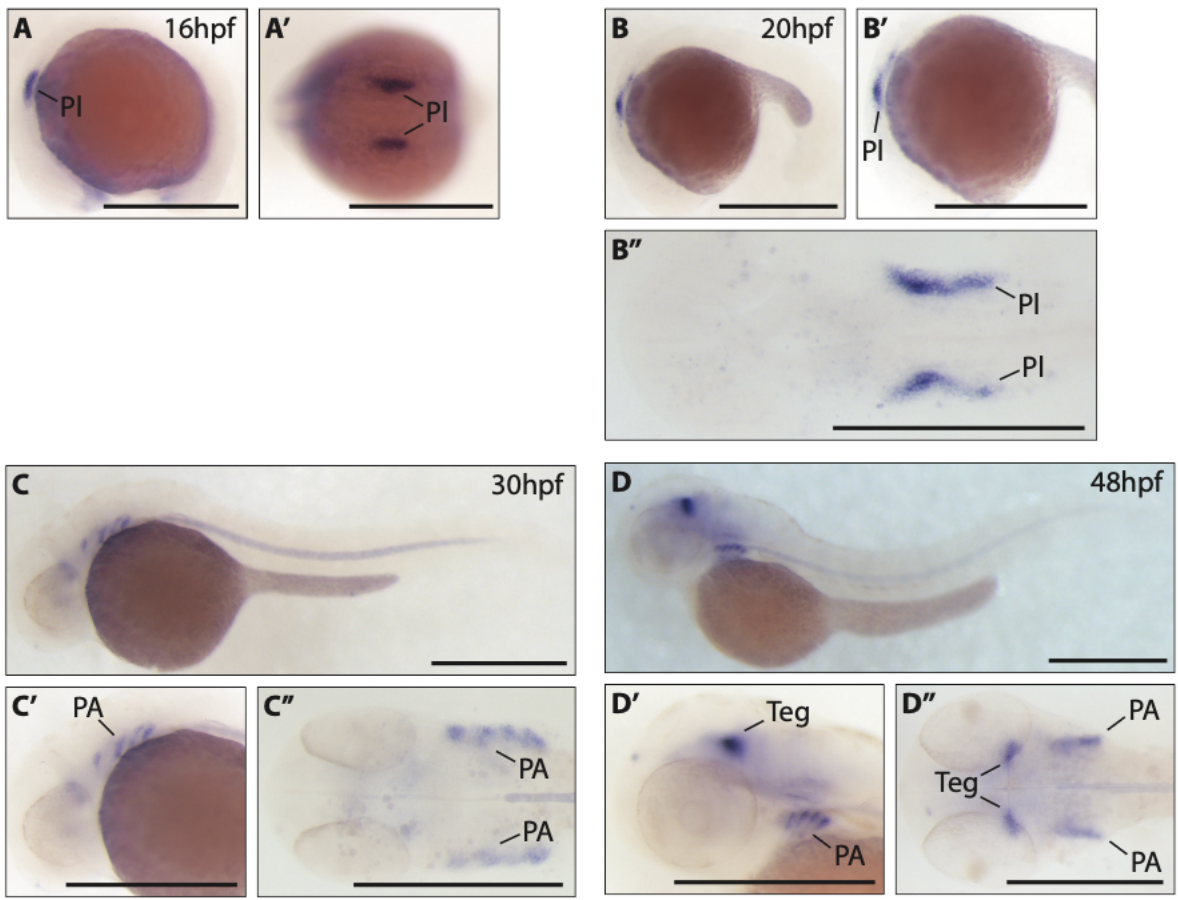

**Figure S11. Expression of *Danio rerio bebe*.** Whole mount *in situ* hybridisation against *bebe* at 16hpf (A, A'), 20 hpf (B-B''), 30 hpf (C-C'') and 48 hpf (D-D''). Lateral whole-embryo views (A-D) and dorsal whole-embryo view (A') are shown. Close-up lateral views (B'-D') and flat-mounted dorsal views (B''-D'') show regions of expression. *bebe* expression is first detected in presumptive placodal cells at 16 hpf, with pharyngeal arch expression at 30 hpf and tegmental expression becoming apparent at 48 hpf. All views oriented anterior to the left. PI, placodes; PA, pharyngeal arches; Teg, tegmentum. Scale bars represent 500 µm.

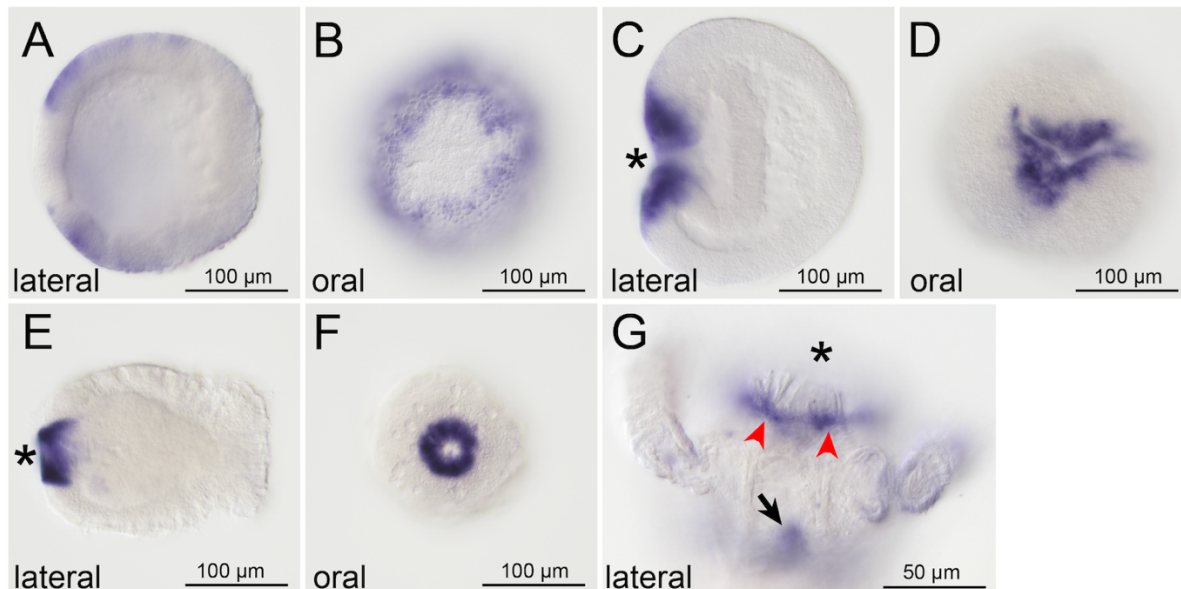

**Figure S12. Expression of *Nematostella vectensis bebe*.** (A-B) Lateral and oral views of a 14 hours post-fertilization blastula stage embryo. *bebe* transcripts are expressed in an oral-to-aboral gradient with a maximum in the future blastopore lip. Oral view shows that the future mesoderm (previously called pre-endodermal plate) is devoid of *bebe* mRNA. (C-D) Lateral and oral views of a 24 hpf mid gastrula. *bebe* transcripts are present only in the blastopore lip. (E-F) Strong *bebe* expression around the mouth of a 4 days post-fertilization planula larva. (G) *bebe* transcripts are expressed (red arrowheads) in a narrow ring of cells surrounding the mouth of the 9 days post-fertilization primary polyp. Black arrow points at the typical unspecific staining in the pre-pharyngeal gastrodermal cells.

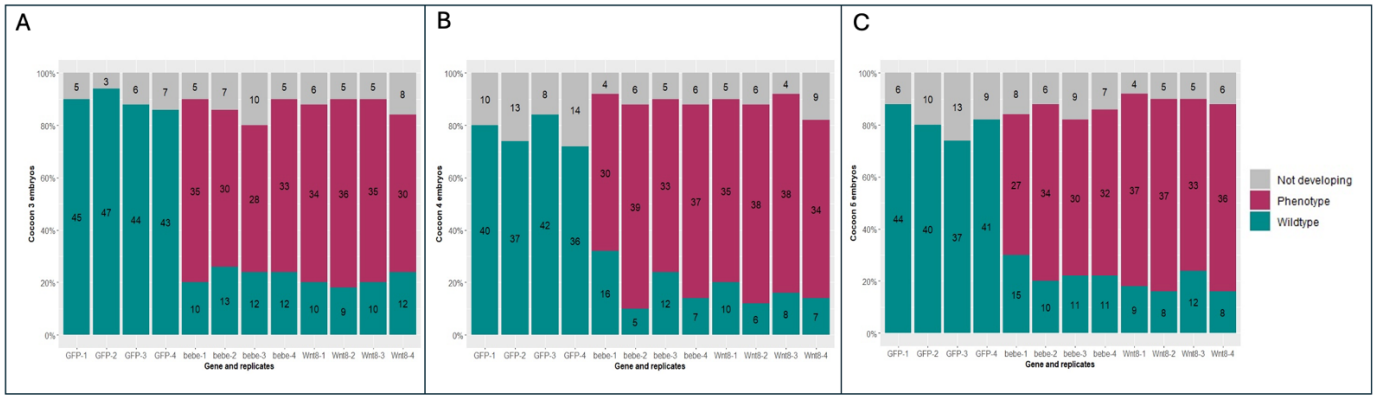

**Figure S13. Frequency of embryonic phenotypes in control and pRNAi cocoons.** Bar charts summarising embryonic phenotypes from double stranded RNA parental injections for GFP (negative control) *Pt-bebe* and *Pt-Wnt8*. A subset of 50 embryos was analysed from 3rd (A), 4th (B), and 5th (C) cocoons from each injected female. 'WT' signifies embryos with normal development (turquoise); 'Phenotype' refers to embryos with mild to extreme knockdown effects (pink); 'Not developing' refers to dead, broken or underdeveloped embryos (grey).

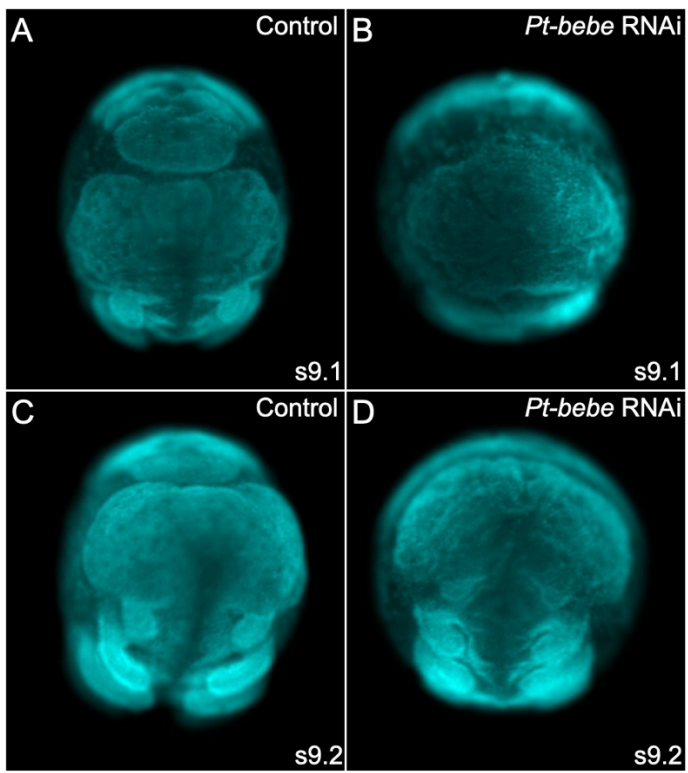

**Figure S14. Developing head morphology of control and *Pt-bebe* pRNAi embryos.** Anterior views showing the head morphology of stage 9.1 and 9.2 control (A and C) and *Pt-bebe* pRNAi knockdown embryos (B and D). Embryos are stained with DAPI and orientated with dorsal up.

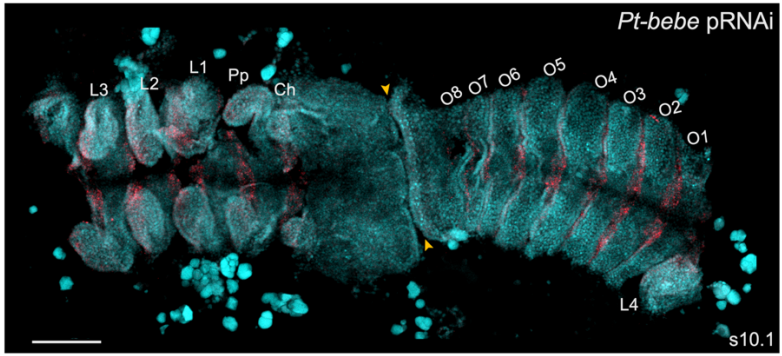

**Figure S15. *Pt-bebe* RNAi embryos do not generate full complement of opisthosomal segments.**

*Pt-bebe* pRNAi embryo showing *Pt-en-1* expression (and DAPI staining) at stage 10.1 with 8 opisthosomal segments. Yellow arrowheads indicate where the distorted SAZ abuts the developing head. Ch, cheliceral segment; Pp, pedipalpal segment; L, leg-bearing segment; O, opisthosomal segment, SAZ, segment addition zone. Flatmounted embryo split between O1 and L4. Scale bar is 200 μm.

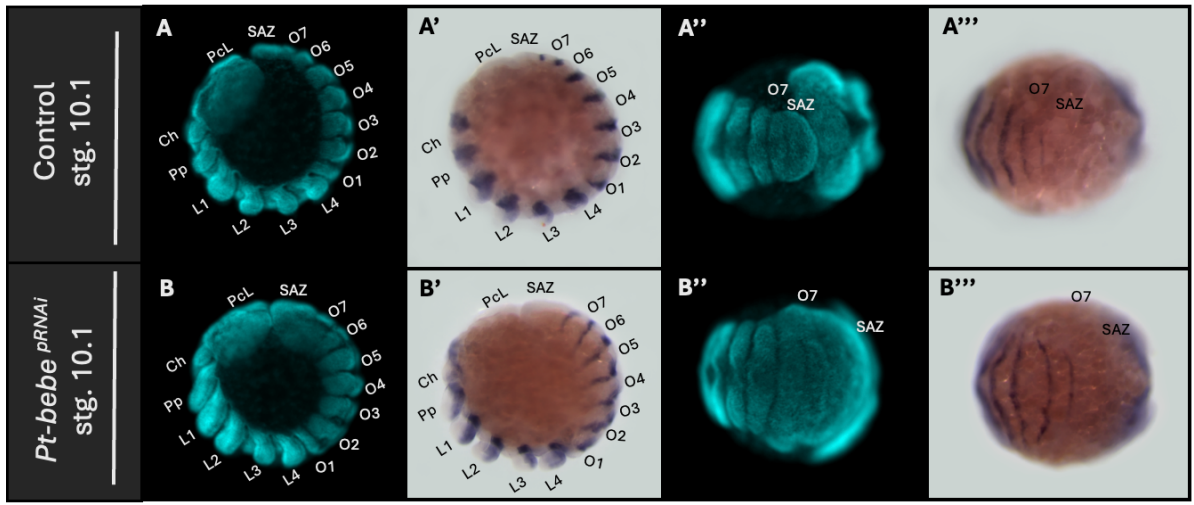

**Figure S16. *Pt-engrailed-1* expression in control and *bebe* RNAi embryos.** *Pt-bebe* pRNAi and control (injected with dsGFP) stage 10.1 whole mount embryos showing *Pt-en-1* expression. Lateral view (left) with DAPI staining (A,B), and *Pt-en-1* expression (A',B'). Posterior ventral view (opisthosoma and SAZ) (A'',A''') and *Pt-en-1* expression (A''', B'''). PcL, pre-cheliceral region; Ch, cheliceral segment; Pp, pedipalpal segment; L, leg-bearing segment; O, opisthosomal segment; SAZ, segment addition zone.

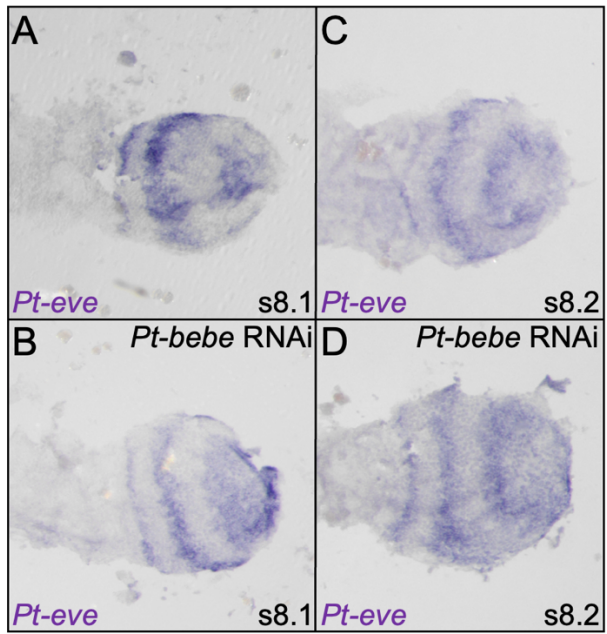

**Figure S17. *Pt-even-skipped* expression in control and *Pt-bebe* pRNAi embryos.**

Expression of *Pt-eve* in control (A, C) and *Pt-bebe* pRNAi embryos (B, D) at stages 8.1 and 8.2.

Embryos are flat mounted with anterior to the left. Only the opisthosomal region and SAZ are shown.

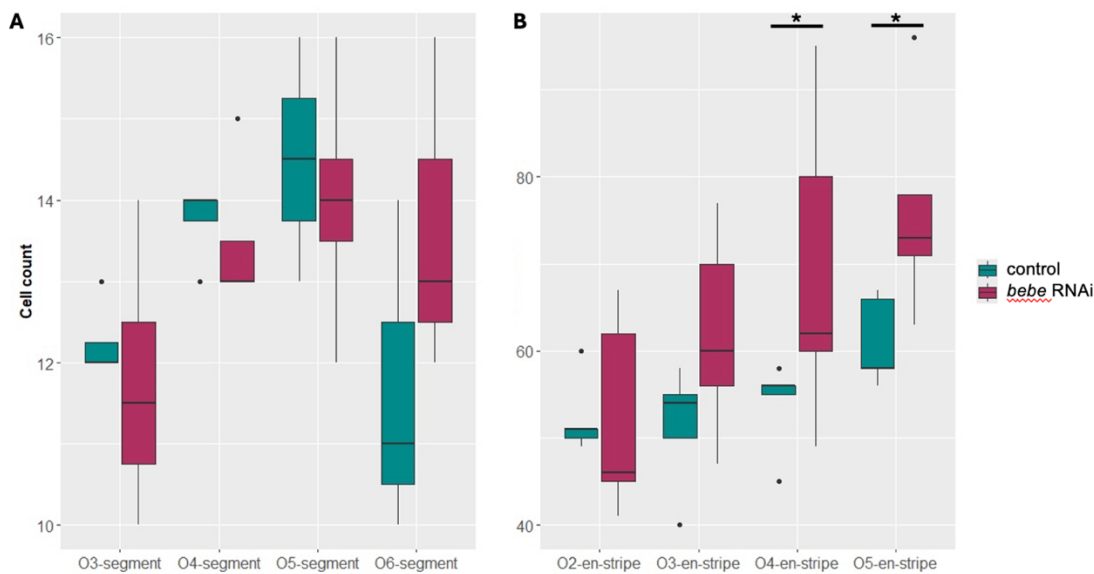

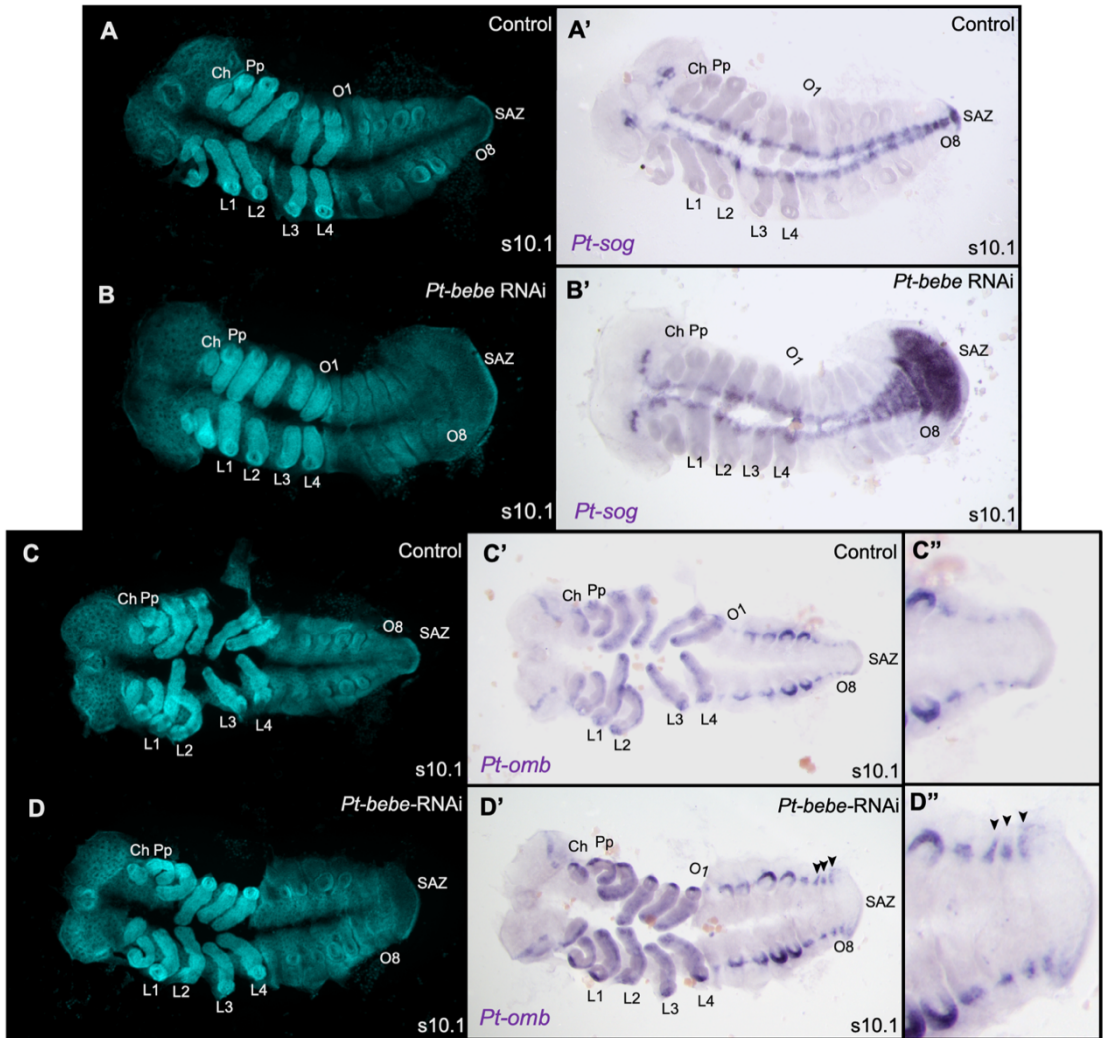

**Figure S19. *Pt-sog* and *Pt-omb* expression after *Pt-bebe* pRNAi knockdown.** Expression of *Pt-sog* in control (A, A') and *Pt-bebe* pRNAi (B, B') embryos at stage 10.1. Expression of *Pt-omb* in control (C-C'') and *Pt-bebe* pRNAi (D-D'') embryos. DAPI stainings (A-D) and brightfield images of *sog* (A' and B') and *omb* (C' and D') expression are shown. Images in C'' and D'' show higher magnification of the posterior opisthosoma and SAZ of the images in C' and D', respectively, with arrowheads indicating extended dorsal stripes of *omb* expression in O6-O8 compared to the control embryo. Embryos are flat mounted with anterior to the left. Ch, cheliceral segment; Pp, pedipalpal segment; L, leg-bearing segment; O, opisthosomal segment; SAZ, segment addition zone.

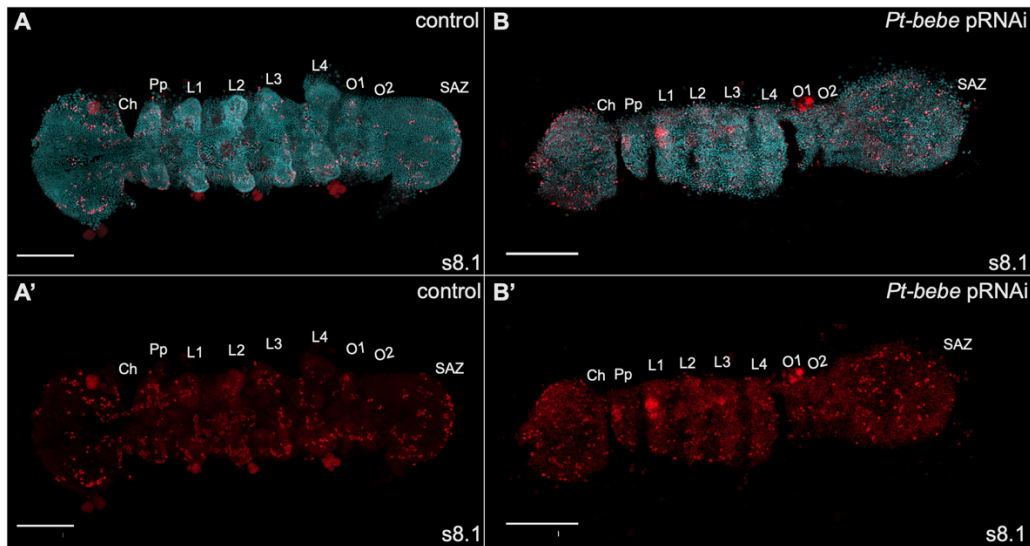

**Figure S20. Dividing cells in control and *Pt-bebe* RNAi embryos.** Anti-Phospho histone H3 staining to indicate dividing cells in control (A and A') and *Pt-bebe* pRNAi knockdown (B' and B') embryos at stage 8.1. Images in A and B show the DAPI staining and those in A' and B' the anti-Phospho histone H3 staining alone. Embryos are flat mounted with anterior to the left. Ch, cheliceral segment; Pp, pedipalpal segment; L, leg-bearing segment; O, opisthosomal segment; SAZ, segment addition zone. Scale bar = 200 μm.

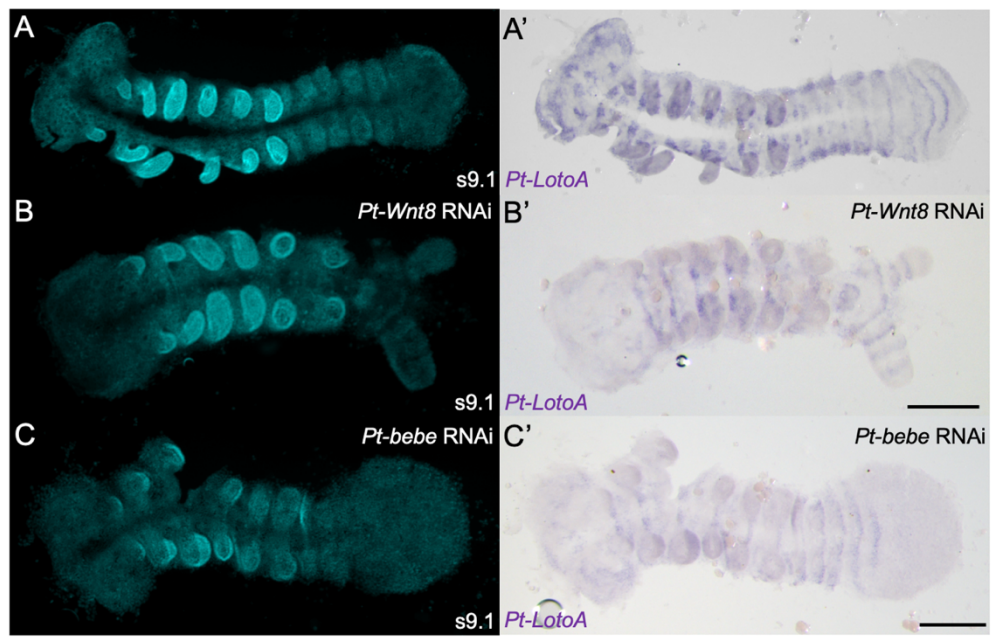

**Figure S21. *Pt-LotoA* expression in control, and *Pt-Wnt8* and *Pt-bebe* pRNAi embryos.** Expression of *Pt-LotoA* in control (A, A'), *Pt-Wnt8* pRNAi (B, B') and *Pt-bebe* pRNAi (C, C') at stages 9.1 and 8.2. Embryos are flat mounted with anterior to the left. DAPI stainings (A-C) and brightfield images of *Pt-LotoA* expression (A'-C') are shown. Scale bars = 200 μm.

### Supplementary File 1. Excel sheet with all Bebe sequence information and primers.
